# Single-Nucleotide-Resolution Computing and Memory in Living Cells

**DOI:** 10.1101/263657

**Authors:** Fahim Farzadfard, Nava Gharaei, Yasutomi Higashikuni, Giyoung Jung, Jicong Cao, Timothy K. Lu

**Author notes:** Corresponding author & lead contact: T.K.L.

## Abstract

Computing and memory in living cells are central to encoding next-generation therapies and studying *in situ* biology, but existing strategies have limited encoding capacity and are challenging to scale. To overcome this bottleneck, we developed a highly scalable, robust and compact platform for encoding logic and memory operations in living bacterial and human cells. This platform, named DOMINO for <u>D</u>NA-based <u>O</u>rdered <u>M</u>emory and <u>I</u>teration <u>N</u>etwork <u>O</u>perator, converts DNA in living cells into an addressable, readable, and writable computation and storage medium via a single-nucleotide resolution read-write head that enables dynamic and highly efficient DNA manipulation. We demonstrate that the order and combination of DNA writing events can be programmed by biological cues and multiple molecular recorders can be coordinated to encode a wide range of order-independent, sequential, and temporal logic and memory operations. Furthermore, we show that these operators can be used to perform both digital and analog computation, and record signaling dynamics and cellular states in a long-term, autonomous, and minimally disruptive fashion. Finally, we show that the platform can be functionalized with gene regulatory modules and interfaced with cellular circuits to continuously monitor cellular phenotypes and engineer gene circuits with artificial learning capacities. We envision that highly scalable, compact, and modular DOMINO operators will lay the foundation for building robust and sophisticated synthetic gene circuits for numerous biotechnological and biomedical applications.

**One Sentence Summary:** A programmable read-write head with single-nucleotide-resolution for genomic DNA enables robust and scalable computing and memory operations in living cells.

## Main Text

Robust and scalable molecular recording and computation platforms in living cells are key to enabling a broad range of bioengineering and biomedical applications. Unlike their silicon-based counterparts that have access to large capacities of addressable memory registers, synthetic genetic circuits currently have very limited information storage capacities and existing methods for encoding information into cellular memory, as well as strategies for integrating such memory with logic operations, are challenging to scale.

Genomic DNA is an ideal medium for biological memory since it is ubiquitously present, naturally replicated at high fidelity within cells, and compatible with natural biological operations. In recent years, several strategies for encoding biological information into DNA and integrating these memories with cellular computers have been described (Farzadfard and Lu, 2014; Kalhor et al., 2017; McKenna et al., 2016; Perli et al., 2016; Roquet et al., 2016; Siuti et al., 2013). However, these methods remain limited in their encoding capacity and scalability. For example, site-specific recombinases that flip or excise targeted DNA segments have been used to create digital memory, sequential logic, and biological state machines in living cells (Roquet et al., 2016; Siuti et al., 2013). However, a different recombinase is required for every unique event that one wishes to record, thus limiting the number of potential states that can be encoded into DNA memory. Furthermore, distances between recombinase-recognition sites usually need to be several hundred base pairs to achieve efficient recombination, thus increasing circuit size (Coppoolse et al., 2005; Stark, 2017). Furthermore, recombinase sites must be pre-engineered into desired target sites, which is time- and labor-intensive, especially if they are to be used in the genomic context.

To address these limitations, we previously developed the SCRIBE DNA writing and molecular recording system, which uses *in vivo* single-stranded DNA expression to generate precise mutations that accumulate into target genomic loci as a function of the magnitude and duration of exposure to an input (Farzadfard and Lu, 2014). However, this approach has been limited to bacteria thus far due to the requirement for specific recombination mechanisms. Alternative molecular recording strategies based on Cas9 nuclease (Kalhor et al., 2017; Perli et al., 2016) have been recently described. However, since the mutational outcomes used to generate memory states by these strategies are generated stochastically, they are not suitable for implementing genetic logic circuits that require robust and deterministic operations. Additionally, due to requirements for host-specific DNA repair and genome editing mechanisms, these systems are only applicable to a subset of organisms.

To overcome these bottlenecks, we describe a platform called DOMINO (for <u>D</u>NA-based <u>O</u>rdered <u>M</u>emory and <u>I</u>teration <u>N</u>etwork <u>O</u>perator) that uses highly efficient and precise DNA writing with CRISPR base editors (Komor et al., 2016; Nishida et al., 2016) to manipulate DNA dynamically and efficiently with single-nucleotide resolution in living cells. DOMINO enables the use of DNA as a uniquely addressable, readable, and writable information storage and computation medium. We demonstrate that the order and combinations of these DNA writing and molecular recording events can be tuned by external inputs and coordinated, allowing one to execute order-independent (e.g., IF EVER A AND IF EVER B), sequential (e.g., A AND THEN B), and temporal (e.g., A AND THEN B after time X) logic and memory operations. DOMINO operators enable highly compact and scalable logic and memory operators that, unlike previous strategies, can be used to realize both digital and analog computation and molecular recording in living cells. Various orthogonal DOMINO operators can be simply created by changing guide RNA (gRNA) sequences, thus making the system highly scalable. These operators can then be layered and interfaced with synthetic or natural regulatory circuits to build more sophisticated genetic programs. Finally, we demonstrate that DOMINO can be combined with established CRISPR-based gene regulation platforms, such as CRISPR interference (CRISPRi) (Qi et al., 2013) and CRISPR activator (CRISPRa) (Farzadfard et al., 2013; Gilbert et al., 2013), to achieve modular and versatile memory and gene regulation programs. DOMINO moves the utility of molecular recording beyond DNA write-only applications – where the recording output can only be read by disruptive DNA sequencing methods – and demonstrates that more advanced biocomputing, such as building logic gates and online state-reporters, can be achieved by coordinating the activity and timing of multiple molecular recorders. These advances address many limitations of the current *in vivo* computing and memory technologies and pave the way towards building advanced genetic circuits with artificial learning capacities.

### Engineering an Efficient Read-Write Head for Genomic DNA

In order to efficiently manipulate genomic DNA in living cells, we used base editing technology (Komor et al., 2016) to build a single-nucleotide resolution “read-write head” for this medium. To this end, we fused Cas9 nickase (nCas9, an addressable DNA “reader” module that is directed by gRNA to bind to specific DNA targets and nicks them) to cytidine deaminase (CDA, a DNA “writer” module that edits the DNA) and uracil DNA glycosylase inhibitor (*ugi*, a peptide which has been shown to improve the DNA writing efficiency by blocking cellular repair machinery) to create CDA-nCas9-ugi. Once localized to the target based on the 12 bp gRNA seed sequence (“READ” address), the writer module can deaminate dC positions in the vicinity of 5’-end of the target (“WRITE” address), thus resulting in DNA lesions that are preferentially repaired as dT (Komor et al., 2016; Nishida et al., 2016). Using cytidine deaminase as the DNA writer module enables dC to dT mutations (or dG to dA mutations if the reverse complement strand is targeted) to be introduced to the WRITE address, resulting in permanent records in DNA. In this memory scheme, an individual mutation or a group of mutations in a target site can be designated as a unique memory state for the corresponding memory register, and mutations introduced by DNA writing events can be considered as transitions between DNA memory states (Fig. 1A). DNA writing events can be controlled by internal or external inputs by placing both the gRNA expression and CDA-nCas9-ugi under regulation by inducible promoters.

**Figure 1.**
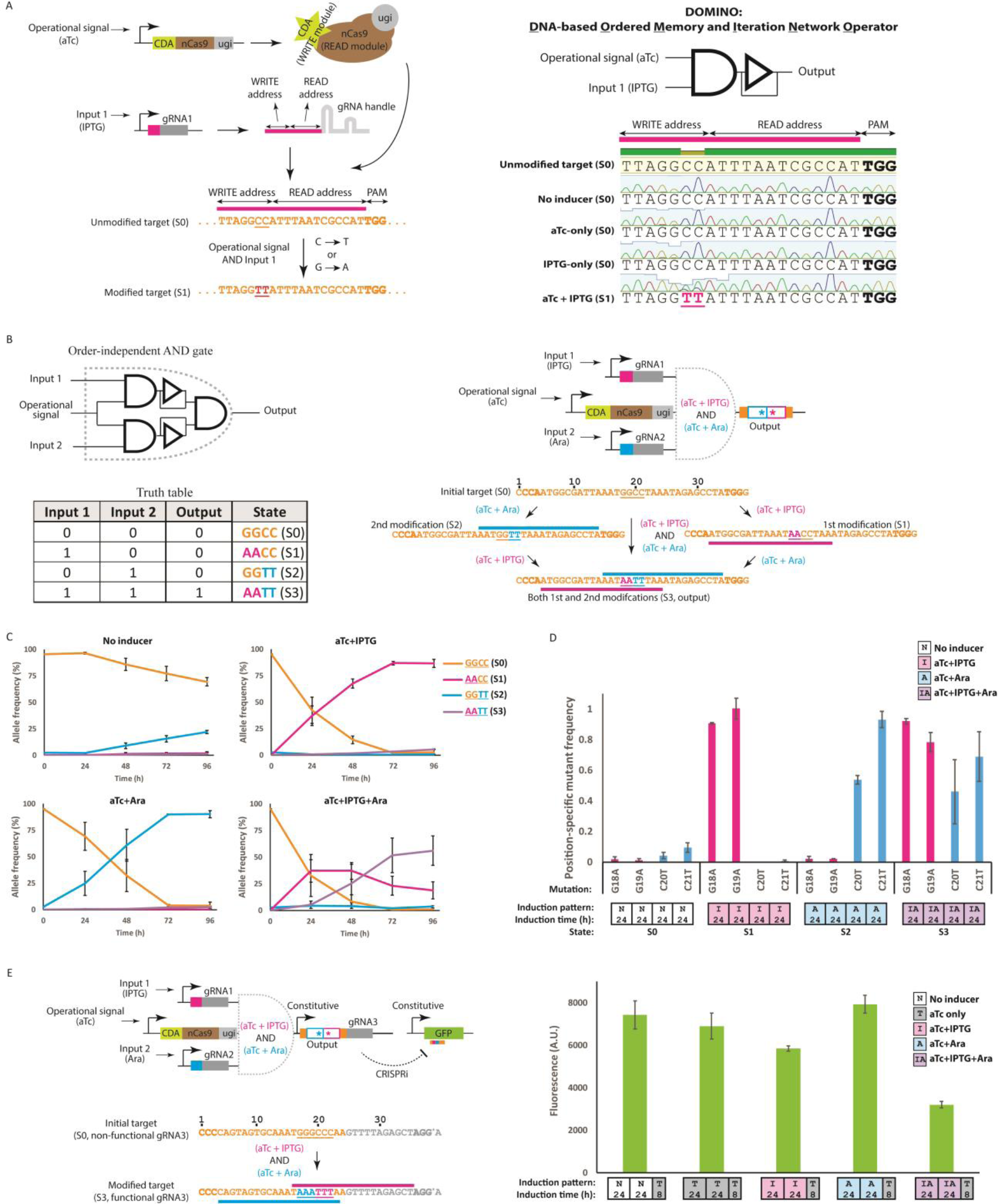
Incorporating memory and logic in living cells by DOMINO. **A)** Schematic representation of DOMINO operators. DOMINO operators are enabled by a DNA read-write head that performs efficient and precise manipulation of genomic DNA with single-nucleotide resolution. In this device, nCas9 (READ module), along with cytidine deaminase (CDA, WRITE module) and uracil DNA glycosylase (*ugi*, WRITE enhancer) domains are addressed to a desired genomic loci using gRNA with a complementary seed region (READ address). Localization of the CDA write module to the target results in the deamination of cytidine (dC) residues in the vicinity of the 5’-end of the gRNA (WRITE address) and their conversion to dU residues, which are then preferentially repaired by the cellular machinery to dT (or dG to dA mutation if the negative strand of DNA is targeted by gRNA). By placing the DNA read-write module and the gRNA under the control of inducible signals, DNA writing for DOMINO operators can be tuned and controlled by external cues. Here, we schematize the basic DOMINO operator as an AND gate since it requires the expression of both the DNA read-write head (i.e., CDA-nCas9-ugi controlled by the “operational signal”) as well as the gRNA (regulated by “Input 1”) with a downstream feedback delay operator (to illustrate the unidirectional and memory aspect of the operator). DOMINO operators can be layered to a wide variety of memory and logic functions. Bold nucleotides on the target show the location of NGG PAM sequence. Targeted nucleotides are underlined. **B)** Order-independent AND gate enabled by DOMINO where the output is ON only when both inputs have been present with any possible order. Induction of the circuit with either of the two inducers (IPTG or Ara), results in editing of the target and transition to an intermediate state (states S1 or S2, respectively). Induction of the circuit with both gRNAs results in generation of the doubly edited DNA sequence (state S3), which is designated as ON state. **C)** Dynamics of allele frequencies obtained by Illumina High-Throughput Sequencing (HTS) for the circuit shown in (B). *E. coli* cells were exposed to different inducer combinations for four days with serial dilution after each 24 hours. Error bars indicate standard deviation of three biological replicates. **D)** Position-specific mutant allele frequencies for the last time point (96 h) of the experiment shown in (C) estimated from Sanger sequencing analysis by Sequalizer (see Supplementary Materials). This data demonstrates the expected outcomes of AND gate behavior at the population level. The x-axis shows dC to dT or dG to dA mutations in the specified positions. For example, the G18A mutation means a dG to dA mutation in position 18 of the target sequence. Small boxes along the x-axis show the induction patterns and duration of induction used in each experiment. For example, the induction pattern of the last sample set ([IA][IA][IA][IA]) means that the samples were induced with aTc + IPTG + Ara for four days with dilutions every 24 hours. Error bars indicate standard deviation of three biological replicates. **E)** The output of DOMINO operators, which is in the form of DNA mutations, can be converted to a gRNA by flanking the target DNA sequence with a desired promoter and gRNA handle. This allows DOMINO operators to be linked to other DOMINO operators or host regulatory networks. To demonstrate this concept, we designed an order-independent DOMINO AND gate with a target sequence flanked by a constitutive promoter and a modified gRNA handle. The modified gRNA handle harbored a dA to dG mutation in a position that was not essential for gRNA function (Briner et al., 2014). This modification (shown by an asterisk) was required to generate an NGG PAM motif for binding of one of the input gRNAs. Upon induction by both inducers, the input gRNAs edit the Specificity-Determining Sequence (SDS) of the output gRNA. The doubly edited output gRNA can then bind to the GFP ORF and repress it via CRISPRi in *E. coli*. In this example, AND logic is realized on the target DNA register (i.e., the output gRNA) while NAND logic is achieved on the output GFP reporter. Error bars indicate standard deviation for three biological replicates.

We demonstrated that this approach enables highly efficient, robust and scalable DNA writing in *E. coli*. We first placed CDA-nCas9-ugi under the control of anhydrotetracycline (aTc)-inducible promoter. Using an Isopropyl β-D-1-thiogalactopyranoside (IPTG)-inducible gRNA as an input, we demonstrated efficient and inducible DNA writing (dC to dT mutations) at desired target sites in the presence of aTc and IPTG induction (Fig. 1A). In this design, which forms the basis of DOMINO operators, the signal controlling the expression of CDA-nCas9-ugi (aTc) that is required for the overall circuit to function can be considered as the “operational signal”, while the signals controlling the expression of individual gRNAs can be considered as independently controllable “inputs”.

### Order-independent DOMINO Logic

DOMINO operators can be arrayed and interconnected in a highly scalable fashion to build robust and complex forms of computing and memory circuits that execute a series of order-independent and/or sequential unidirectional DNA writing events. The frequency and order of these DNA writing events can be controlled by internal and external cues, as well as by carefully selecting the position of mutable residues within the target. For example, by layering two DOMINO operators, we built a two-input order-independent AND logic gate, where the A AND B logic is executed independent of the order of addition of the inputs (Fig. 1B). In this design, two distinct gRNAs were placed under the control of IPTG- and Arabinose (Ara)-inducible promoters, respectively. In the presence of its corresponding inducer, each gRNA is expressed and directs the DNA read-write module (which itself is expressed in the presence of the operational signal, aTc) to its cognate target site, resulting in precise dC to dT mutations (or dG to dA mutations in cases where the gRNA targets the reverse-complement strand) within the WRITE address.

To assess the performance of the order-independent DOMINO AND gate, we induced cells harboring this circuit with different combinations of the inducers for multiple days and analyzed dynamics of allele frequencies at the target locus by high-throughput sequencing (HTS) over multiple time points. As shown in Fig. 1C, in the presence of the operational signal (aTc) and each of the two inputs (IPTG or Ara), mutations were accumulated in the target sites of the induced gRNA in a linear fashion within the population and comprised ∼100% of the population after 72 hours of induction. This corresponds to transitions from the unmodified state (state S0) to either of the two singly modified states (state S1 or S2). The time required for transitioning between the two states can be considered as the “propagation delay” of the corresponding DOMINO operator. On the other hand, when cells were induced with both inputs (IPTG AND Ara), the target sites for both gRNAs were edited, resulting in the accumulation of doubly edited sites (state S3) in the target locus. We defined states S0, S1, and S2 as the OFF states and S3 as the ON state, which means that this system implements AND logic. In this experiment, low levels of a singly mutated allele (state S2) accumulated in the absence of any induction, likely due to leakiness of the Ara-inducible promoter (pBAD) in these cells and/or high binding efficiency of its corresponding gRNA. The performance of the circuit should be improved by lowering the Ara-inducible promoter’s basal activity, for example, by overexpressing pBAD repressor (*araC*) or using tighter promoters (Arpino et al., 2013), or alternatively, by lowering the copy numbers of DOMINO operators (Lee et al., 2016). Nevertheless, the doubly edited allele (state S3) only accumulated in the presence of both IPTG and Ara, indicating the robust AND logic can be achieved despite the leakiness of one of the input promoters.

Notably, these results show that in DOMINO operators, the accumulation of the singly mutated alleles in the presence of the operational signal and individual inducer inputs follows a linear trend over the course of few days. About 3 days was required for the unmodified allele to be fully converted into the modified allele(s), thus indicating the propagation delays of the corresponding operators. This feature enables one to use DOMINO to implement both analog and digital computing, since continuous changes that occur within the propagation delay window can be used to implement analog computation, while fully converted states can be considered as transitions between digital states and thus used for digital computation.

The states designated in the AND gate logic described in this example are arbitrary defined; for example, the doubly mutated allele (state 3) was defined as the ON state. The same circuit can be defined, for example, as a NAND gate if the unmodified state (state S0) is designated as ON (“1”) output and states S1 through S3 are designated as OFF (“0”) outputs. Alternatively, each of the four different mutational states can be defined as distinct outputs, in which case the circuit can be considered as a 2-input/4-output decoder.

In this experiment, two mutable residues within the editing window of each gRNA were used, and the memory states were defined so that mutations in both of these residues were required to be considered as a state transition. One could define mutations in only one of the two nucleotides available for editing as intermediate states (that can be discarded), or if desired, as usable transient memory states. Furthermore, the number of memory states as well as the response dynamics (e.g., propagation delay) for each DOMINO operator can be tuned by using different numbers of mutable residues (dC or dG) within the WRITE window, or adjusting the position of these residues within this window.

While HTS offers a powerful way to quantify the outcome of DOMINO circuits, its relatively high cost inspired us to develop a strategy for using Sanger sequencing chromatograms to quantify position-specific mutant frequencies within a mixture of DNA species. This algorithm, named Sequalizer (for <u>Sequ</u>ence equ<u>alizer</u>), normalizes Sanger chromatogram signals and calculates the difference between the normalized signals from a test sample and an unmodified reference to identify position-specific mutations. It then uses this calculated difference to estimate position-specific mutant frequencies at any given target position. We validated the accuracy of this method by constructing a standard curve based on known ratios of mutant and wild-type (WT) sequences, and comparing the Sequalizer results with next-generation sequencing (see Supplementary Materials and Fig. S1). The Sequalizer output, which is based on population-averaged Sanger sequencing results, provides an estimate of position-specific mutant frequencies in an entire population. Though Sequalizer does not always provide accurate absolute values of mutant frequencies, fold changes in estimated mutant frequencies are accurate (see Supplementary Materials and Fig. S1C). Additionally, unlike HTS, Sequalizer output does not provide insights into the identities and frequencies of individual alleles in the population. Nevertheless, given the high specificity of the DNA writers and predefined target sites for DNA writing, this approach can be used as a low-cost alternative to HTS to assess performance of DOMINO and other precise genome-editing platforms.

In addition to HTS, we analyzed the samples obtained from experiment shown in Fig. 1B by Sanger sequencing and Sequalizer. As shown in Fig. 1D and Fig. S1C, in samples induced with either of the two inputs, the frequencies of mutants in positions corresponding to the cognate target sites of the induced gRNA increased in the population. On the other hand, in samples that were induced with both gRNAs, the mutation frequencies in the target sites of both gRNAs were increased (state S3). These results demonstrate that Sequalizer results are consistent with and could accurately estimate changes in position-specific mutant frequencies obtained by HTS.

In addition to AND gate, other logic can be readily implemented by carefully positioning mutable residues on the targets, as well as designing the combinations and order of DNA writing events. Furthermore, additional input gRNAs can be incorporated to achieve operators with more than two inputs, thus demonstrating scalability of this approach (Fig. S2).

The output of DOMINO operators takes the form of DNA mutations that accumulate at a target site. One can flank this target site with a desired promoter and a gRNA handle to convert the output of a given DOMINO operator into downstream gRNA expression. The output gRNA can then be interconnected with other DOMINO operators to build more complex circuits. In addition, it can be combined with CRISPR-based gene regulation platforms such as CRISPRi and CRISPRa to dynamically regulate cellular phenotypes. To demonstrate this, we engineered an AND operator by layering two DOMINO operators under the control of inducible promoters to edit a third gRNA as the output (Fig. 1E). The input gRNAs were controlled by IPTG- and Ara-inducible promoters, respectively. In the presence of both inducers, the output gRNA was modified by both input gRNAs such that it could then bind to and repress a downstream reporter gene (GFP) (Fig. 1E, aTc + IPTG + Ara co-induction for two 24-hour periods followed by aTc-induction for 8 hours: [IA][IA][T] induction pattern). When targeting gRNA as an output, both the Specificity Determining Sequence (SDS) of the output gRNA as well as its constant region (handle) can be modified. Mutating the SDS is useful when the creation of a unique gRNA is the desired output. On the other hand, mutating the gRNA handle enables one to activate/deactivate an entire set of gRNAs. Furthermore, one can also target gene regulatory and functional elements, such as promoters, ribosome binding sites, start/stop codons, as well as active sites within proteins to tune the expression or activity of downstream components as shown in Fig. S3.

### Sequential DOMINO Logic

In addition to realizing order-independent logic, one can carefully control the sequence and timing of DNA writing events executed by DOMINO operators to achieve sequential logic, where desired outputs are generated only when the correct order of inducers is added. To achieve this, for example, one can design the gRNA output of one operator to be used as the input for a downstream operator (Fig. S2C). This design can be used to functionally connect DOMINO operators that are not physically co-located, and offers control over the individual DOMINO operators. Alternatively, sequential logic can be achieved by overlapping mutable residues in the WRITE address of one operator with the READ address of a downstream operator (Fig. 2). This design uses DNA mutations rather than cascades of gRNAs as a way to interconnect *cis*-encoded DOMINO operators, thus offering a highly compact and scalable strategy for encoding sequential logic.

**Figure 2.**
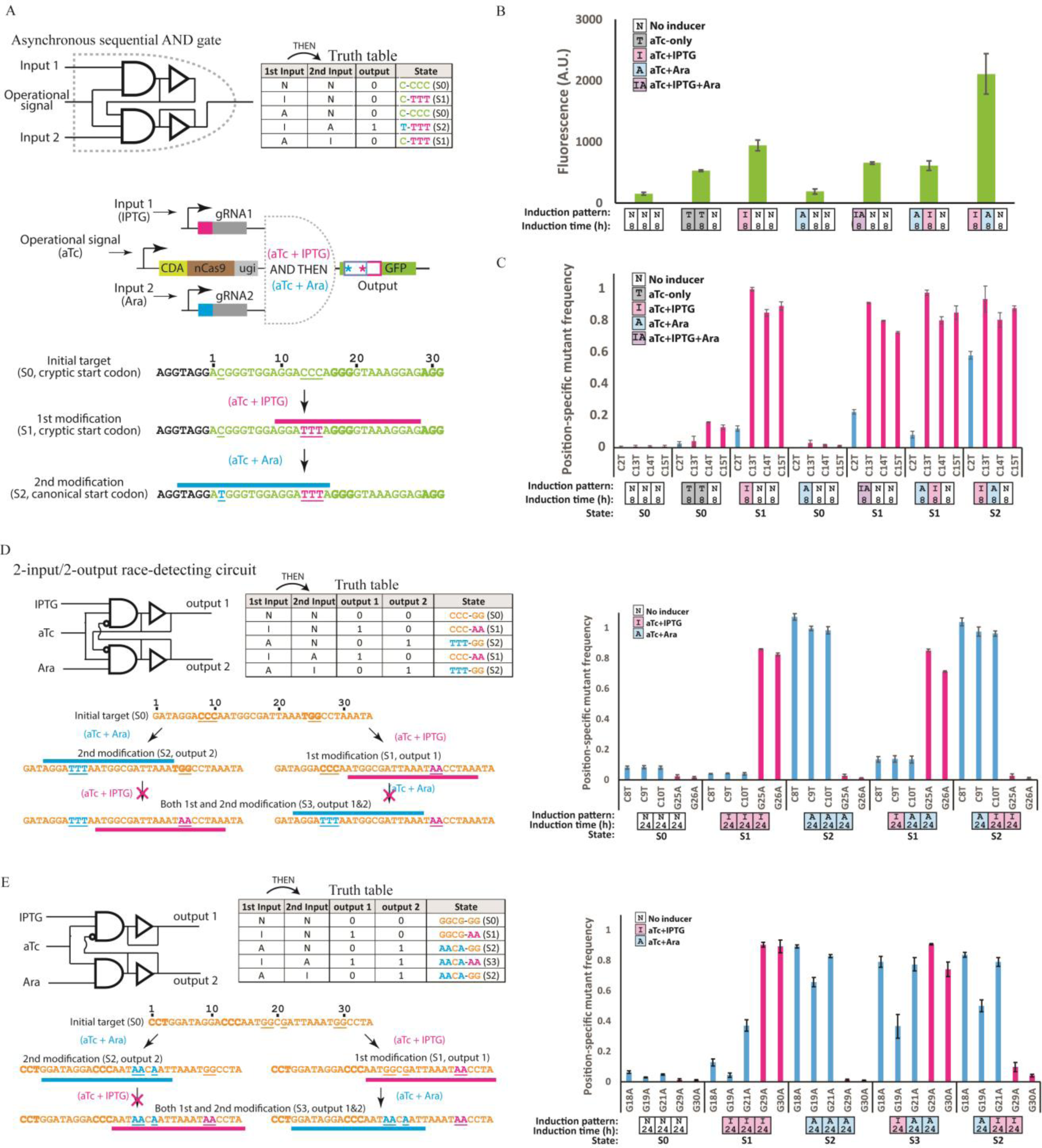
Building sequential logic by DOMINO operators. **A)** Sequential AND gate encoded with DOMINO operators. The output of a DOMINO operator was used as an input for another operator, which in turn mutates a non-canonical start codon (ACG) within the GFP ORF into a canonical (efficient) start codon (ATG), thus increasing GFP signal. The second gRNA (induced by Ara) can bind to and enact the start-codon mutation only after the first gRNA (induced by IPTG) has edited its target. **B)** GFP signal measured by flow cytometry for the circuit shown in (A). Only when IPTG AND THEN Ara were applied was the sequential logic satisfied, thus resulting in increased GFP signal. Error bars indicate standard deviation of three biological replicates. **C)** Position-specific mutation frequency obtained from Sequalizer analysis for the experiment shown in (A). Consistent with GFP data, the highest frequency of ACG to ATG conversion (blue bars) was achieved when the samples were induced with IPTG AND THEN Ara. Error bars indicate standard deviation for three biological replicates. **D)** Two-input/two-output race-detecting circuit. Two gRNAs were designed so that editing by one gRNA destroys the PAM domain for the other gRNA, thus inhibiting its binding. Sequential expression of each gRNA resulted in an output corresponding to the output of the first gRNA, independent of whether the second gRNA was expressed or not. Error bars indicate standard deviation for three biological replicates. **D)** Another example of sequential DOMINO logic, where sequential induction of cells with IPTG AND THEN Ara results in the sequential transition between two modified states (states S1 and S3, respectively). However, induction of cells with the reverse order (Ara AND THEN IPTG) only results in a one-step transition to state S2. Error bars indicate standard deviation for three biological replicates.

To demonstrate the latter strategy, we first constructed an asynchronous sequential AND gate, where sequential addition of the two inputs in the correct order (IPTG AND THEN Ara) leads to mutation of a cryptic start codon (ACG) into the canonical (and more efficient) start codon (ATG) in the GFP ORF, thus increasing the GFP signal (Figs. 2A and 2B). We observed slight increases in GFP signal in cells that had been induced with the first inducer (i.e., IPTG) or those that had been co-induced with both inducers (Fig. 2B). The former was likely caused by the leakiness of the second (Ara-inducible) promoter while the latter was likely due to the simultaneous presence of both inducers in the media, which could result in the execution of sequential DNA mutations in the correct order to some extent. Nevertheless, the GFP signal was significantly higher when cells were exposed to the correct order of the inducers. We further confirmed these results by analyzing Sanger sequencing chromatograms by Sequalizer (Fig. 2C). Consistent with flow cytometry data, samples induced with the correct order of the inputs showed the highest level of the dC to dT mutation in the position corresponding to the cryptic start codon (Fig. 2C), indicating the execution of a cascade of DNA writing events that lead to execution of sequential AND logic. 9

As another example, we built an asynchronous 2-input/2-output race-detecting circuit, where the output of the circuit is determined by the inducer added first and not the other inducer added second (Fig. 2D). In this design, the PAM domain for each gRNA is placed within the WRITE window of the other, in a way that editing mediated by one gRNA destroys the PAM domain for the other gRNA, thus preventing binding and subsequent editing by that gRNA. As shown in Fig. 2D, Sequalizer analysis of cells induced with different combinations of inducers showed that the output of the circuit depends on the identity of the first inducer. Specifically, cells that were first induced with IPTG were converted to state S1, independent of addition of the second inducer (Ara) at a later stage, and those cells that were first induced with Ara were converted to state S2 independent of IPTG induction.

When cells were induced with IPTG AND THEN Ara (Fig. 2D, IPTG induction for one day AND THEN Ara induction for two days ([I][A][A] induction pattern)), we observed a slight increase in the mutant frequency in the positions corresponding to targets of the Ara-inducible gRNA. We suspected this to be due to leakiness of the Ara-inducible promoter during IPTG induction period (i.e., before ending the propagation delay of the first operator), which would lead to expression of gRNA2 and aberrant transition of a small subpopulation of cells to state S2. Nevertheless, since editing by one gRNA should destroy the PAM domain for the second gRNA, the race-detecting logic should still hold within each single DNA molecule. High-throughput sequencing of these samples revealed that indeed this was the case since doubly edited allele (i.e., state S3, corresponding to editing events by both gRNAs) were extremely rare (Fig. S4A).

This experiment indicates that the ratio between edited alleles in a population can be tuned by controlling the induction time of each of the inputs, while ensuring that the desired logic is applied at the level of each individual DNA molecule. Alternatively, if conversion of the whole population to a final state is desired, one can perform each induction step for periods longer than operator’s propagation delay (i.e., multiple days) to allow the full conversion of cells to a given state before moving to the next induction step. This control over the degree of commitment of cells to different states could be useful for dividing biological tasks between different subpopulations in a community. For example, one subpopulation of cells could be edited to activate metabolic pathway 1 and the other subpopulation of cells could be edited activate metabolic pathway 2; the relative ratio of activation could be tuned using our DOMINO circuits to control the overall population performance. 10

Finally, we constructed a 2-input/2-output sequential logic circuit, where induction with IPTG AND THEN Ara results in step-wise transition between two modified states (a sequential AND gate) while induction in the opposite direction (i.e., Ara AND THEN IPTG) results in transition to a different state. In this circuit, editing mediated by one gRNA destroys the binding site of the other gRNA, while editing mediated by the second gRNA does not interfere with the binding or editing of the first gRNA. As shown in Fig. 2E, this circuit is an intermediate circuit between the sequential AND gate (Fig. 2A) and the race-detecting circuit (Fig. 2D). Induction of this circuit with IPTG resulted in the transition of the target register from the initial unmodified state (state S0) to the first modified state (state S1). Subsequent induction of these cells with the second inducer (Ara) led to transition of these cells to the doubly mutated state (state S3). On the other hand, when cells were first induced with Ara, they were converted to an alternative singly modified state (state S2). However, subsequent induction of these cells with IPTG did not result in a transition, thus realizing the expected behavior. Using high-throughput sequencing, we confirmed that expected transitions between the states, and thus the circuit logic, held at the single-molecule level (Fig. S4B).

### Temporal DOMINO Logic

The above examples demonstrate that the sequence and timing of DNA writing events mediated by DOMINO operators can be controlled by external cues. In addition to building sequential logic, where the execution of events in a specified order leads to a desired output, the propagation delay in DOMINO operators can be exploited to incorporate temporal logic into circuits, where a desired output is produced only after a certain period of time has passed. In a simple form, DOMINO delay operators can be built by constructing a series of overlapping repeats to act as target sites for a desired gRNA (Fig. 3A). This repeat configuration allows one to overlap the READ address of each gRNA operator site with the WRITE address of the previous gRNA. Initially, the gRNA can bind to the first (i.e., 3’-end) repeat, but not to the upstream copies of the repeat that harbor dC residues (instead of dT) in the sequence corresponding to the gRNA READ address (i.e., the gRNA seed sequence). Upon binding to the first repeat, the gRNA can mutate the dC residues in the repeat immediately upstream of its binding site (i.e., the second repeat), thus converting that repeat to a new binding site for another copy of the same gRNA. This process is sequentially repeated to generate new binding sites for the gRNA. Much like an array of physical domino pieces that fall down one by one, each genome-editing event is initiated only after editing in the previous repeat has occurred, thus ensuring a sequential cascade of DNA writing events. The total delay can be tuned by changing the number of the repeats, modifying the overlapping distance between the repeats, or adjusting the distance of mutable residues from their corresponding PAM sequences.

**Figure 3.**
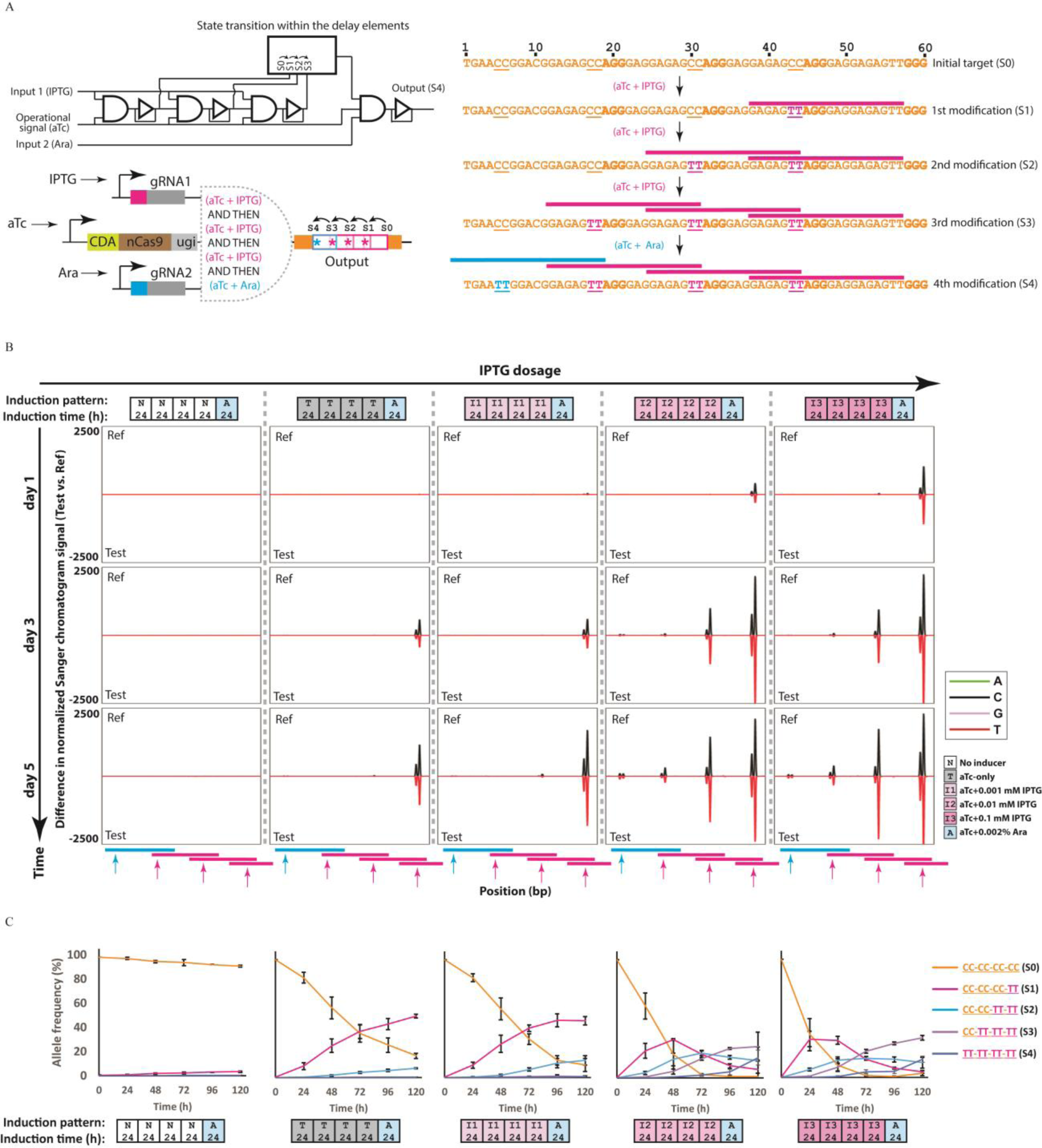
Incorporating propagation delay and temporal logic into living cells. **A)** Time-dependent logic and tunable propagation delay can be programmed by DOMINO operator cascades. DOMINO operators possess an inherent propagation delay (the time required for transition from a non-modified state to modified state) that can be modulated in an analog fashion (stronger induction results in a shorter delay). Multiple DOMINO operators can be placed sequentially in an array to build longer delays and then coupled with other logic operators to build temporal logic. We constructed a series of overlapping repeats to serve as gRNA binding sites. Once expressed, the first gRNA (IPTG-inducible, pink) can bind to the downstream repeat, but not to the other instances of the repeats due to presence of dC residues in these repeats that form mismatches with the gRNA READ address. Upon binding the downstream repeat, the DNA read-write head can mutate these dC residues to dT in the immediately adjacent upstream repeat, thus creating a new binding site for this gRNA. In turn, this event recruits the read-write head once again and makes the third repeat available for binding. The second gRNA, which is under control of Ara, is only able to bind to and edit its target when the third copy of the repeat is edited by the first gRNA, thus encoding time-dependent sequential logic. **B)** *E. coli* cells harboring the circuit shown in (A) were exposed to different concentrations of the first inducer (IPTG) for 4 days with serial dilution after each day, followed by a one-day exposure to the second inducer (Ara). The propagation of the signal as manifested by sequential mutations in the repeat array was monitored by analyzing Sanger chromatograms with Sequalizer. Transitions between states occurred in a time- and IPTG-dosage dependent fashion, and only cells exposed to higher concentrations of IPTG (0.1 mM and 0.01 mM) accumulated mutations to the level that enabled a response to the second inducer (Ara) by the last day of experiment. **C)** Transitions between the memory states for samples shown in (B) assessed by HTS. Error bars indicates standard deviation for three biological replicates.

In addition, the output of the delay elements can be combined with additional logic operators and internal or external cues to create more complex forms of temporal logic. To demonstrate this concept, we placed three DOMINO delay elements into an array and linked the output of the array to a second DOMINO operator that implements sequential AND logic (Fig. 3A). This design achieves temporal and sequential AND logic since the first (IPTG-inducible) gRNA has to execute three consecutive DNA writing events before the Ara-inducible gRNA corresponding to the last operator can bind to and edit its target. We induced cells harboring this circuit with different IPTG concentrations for 4 consecutive days followed by a final day of induction with Ara. Using Sanger sequencing on the population and Sequalizer analysis, we observed a time- and IPTG-dosage-dependent accumulation of mutations in the target sites within repeats, corresponding to propagation of the signal through the repeat array (Fig. 3B). The rate of propagation of the mutation cascade through the delay elements correlated with both the concentration and duration of exposure to IPTG. By the end of the experiment, mutations in the position corresponding to the target site of the second gRNA (shown by the blue arrow in Fig. 3B) were detected only in conditions in which mutations had accumulated through the entire cascade, corresponding to the samples that had been induced with the highest IPTG concentrations.

We further confirmed these results by analyzing these samples with HTS. This analysis also showed time- and IPTG-dosage-dependent mutation accumulation within the repeats (Fig. 3C). Furthermore, the mutation corresponding to the target of the Ara-inducible gRNA only accumulated in the later time points and only in cultures induced with high concentrations of IPTG. Upon induction of the samples by Ara, the frequency of the allele corresponding to the final output of the circuit (i.e., state S4) only increased in samples that had been previously induced with high IPTG concentrations (i.e., 0.01 mM and 0.1 mM). These results further demonstrate that, in addition to enacting delays in gene circuits, an array of DOMINO delay elements can be used as a multi-state memory register that undergoes transitions between different discrete states (i.e., sequential mutations) in a time- and dosage-dependent fashion. In this design, the number of memory states can be tuned by changing the number of repeats. Moreover, the timing and probability of transitions between repeats can be adjusted by changing the position of mutable residues within the repeat overlaps, or tuned dynamically by external cues.

Finally, to demonstrate the power of the technique, we used DOMINO delay elements to build a gene expression program in which the conversion of cryptic ACG start codons into canonical ATG start codons in three different ORFs was temporally controlled by a single input (Fig. S5). We envision that more complex versions of temporal logic, such as counters, can be constructed by integrating delay elements into multiple-input DOMINO operators.

### Associative Learning Circuits and Online DNA-State Reporters

A unique feature of DOMINO operators compared to other memory platforms is that the DOMINO DNA read-write head can be further functionalized with additional effector domains, such as transcriptional activators and repressors, to achieve combined DNA writing and transcriptional regulation. This offers the unprecedented capacity to perform both genetic and epigenetic modulation and thus combine DNA memory states with functional outcomes. For example, this feature enables the construction of circuits that can learn and remember. Specifically, we devised a synthetic gene circuit that undergoes associative learning (Bray, 2003; Gandhi et al., 2007; Nesbeth et al., 2016; Tagkopoulos et al., 2008) such that its gene expression output is reinforced by a given stimulus (Figure 4A). While transcriptional positive feedback loop can also be used to implement synthetic self-reinforcing circuits, the state of such circuits can fluctuate due to their reliance on continuous transcription for state maintenance. In contrast, an associative learning circuit that uses genetically encoded memory to gradually reinforce a response remains intact and stable even after the initial stimuli is removed.

**Figure 4.**
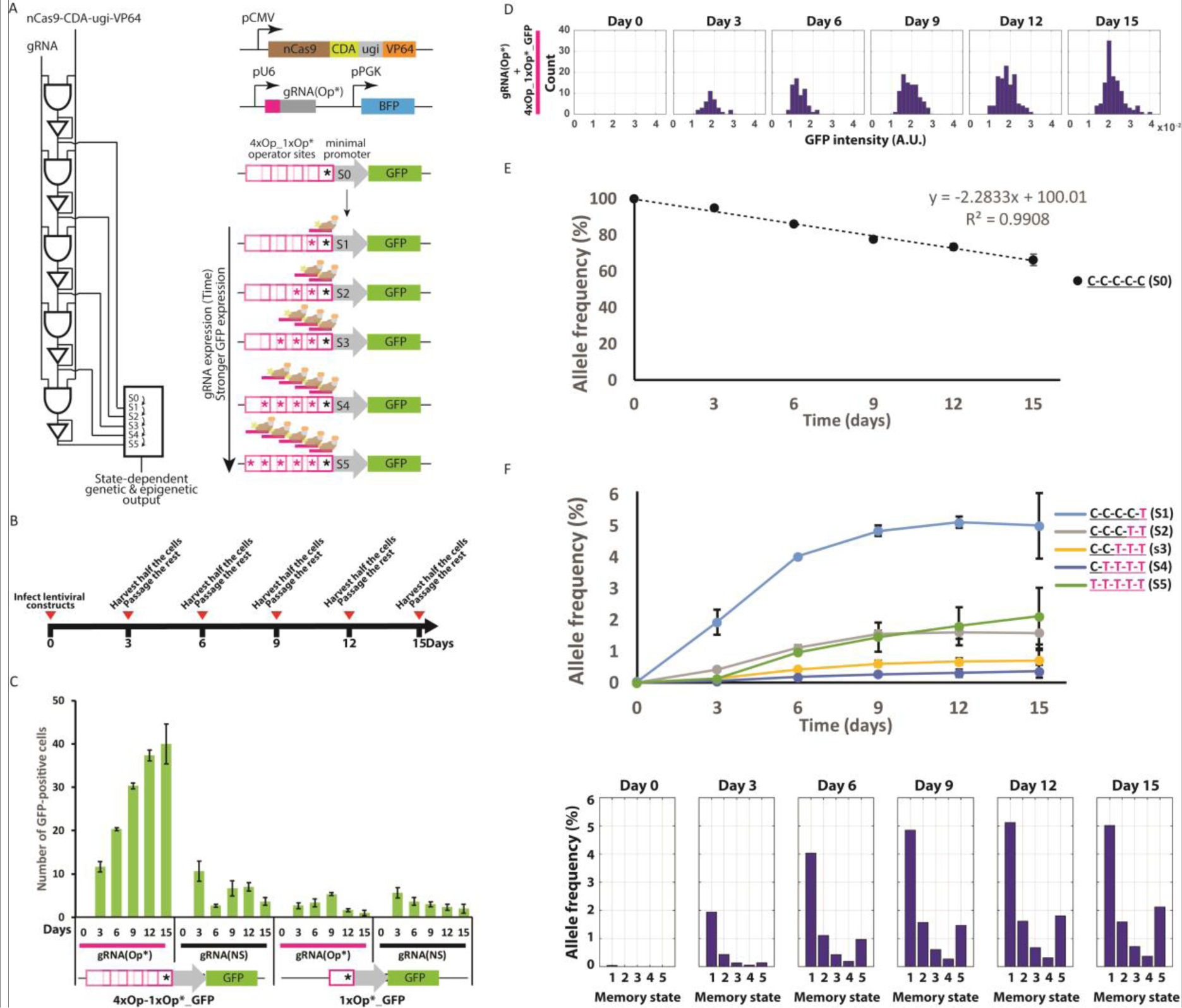
Associative learning and online DNA-state reporting circuits in human cells. **A)** Being CRISPR-Cas9-based, DOMINO operators can be functionalized with transcriptional and epigenetic modules to implement gene regulation integrated with computing and memory. As an example, we functionalized the read-write head with a transcriptional activator (VP64) and used it to sequentially edit and activate multiple operator sites that were arrayed in overlapping repeats (composed of four copies WT unmutated repeats (Op) followed by a downstream mutated repeat (Op*)) upstream of a minimal promoter (4xOp_1xOp*_GFP). In the presence of an Op*-specific gRNA (gRNA(Op*)), this system allows for sequential conversion of Op sites to Op* and binding of the transactivator to the progressively mutated operator sites in the promoter, which in turn results in GFP signal increases. Therefore, cells harboring this circuit manifest sequential and permanent transitions between DNA states and increases in GFP in response to increased gRNA expression over time. Thus, the circuit can be considered as an example of associative learning. **B)** HEK 293T cells were transfected with the circuit shown in (A) via a two-step lentiviral delivery protocol and were grown with serial passaging every three days as indicated. At the end of each passage, GFP signal was assessed by microscopy and DNA memory state was assessed by HTS. **C)** The mean number of GFP-positive cells in different samples harboring either the Op*-specific gRNA (gRNA(Op*)) or a non-specific gRNA (gRNA(NS)) and either 4xOp_1xOp*_GFP or 1xOp*_GFP as the reporter. The number of GFP-positive cells harboring 4xOp_1xOp*_GFP and gRNA(Op*) increased over time. In contrast, the number of GFP-positive cells in cultures harboring gRNA(NS) or 1xOp*_GFP and gRNA(Op*) did not change and remained at background levels. **D)** Histogram of signal intensities for GFP-positive cells harboring 4xOp_1xOP*_GFP and gRNA(Op*). The gradual increase in GFP signal intensities is reflected as a shift to the right in the histograms, indicating multi-stage GFP activation in these cells. **E)** Dynamics of the frequency of the WT unmodified allele (state S0) in cultures harboring 4xOp_1xOp*_GFP and gRNA(Op*) assessed by HTS. The frequency of the unedited allele decreased linearly over time, indicating that the DNA writing circuit can be used as an analog recorder for the input gRNA. **F)** Dynamics of mutant allele frequencies (memory states S1 through S5) for the same samples as (E), shown as time-series data and histograms. Consistent with the GFP data, the first four memory states (S1 through S4) started to accumulate sequentially (state S1, then state S2, then S3 and then S4) until they reached a plateau. Moreover, memory state S5, which corresponds to the highest GFP expression state, increased steadily over time, as is expected from the terminal product of the DNA memory circuit.

To demonstrate this concept, we first made an array of overlapping repeats (operators) composed of four WT repeats (4xOp) and a downstream mutant repeat (1xOp*) which harbored a dC to dT mutation. We then placed this repeat array upstream of a minimal promoter driving GFP to build the 4xOp_1xOp*_GFP reporter construct. Additionally, we built a second reporter (1xOp*_GFP) by placing a single Op* repeat upstream of the minimal promoter driving GFP. We also functionalized the DNA read-write head (nCas9-CDA-ugi) with a transcriptional activator domain (VP64) and cloned the nCas9-CDA-ugi-VP64 fusion construct along with either of the two reporter constructs into lentiviral vectors, which were subsequently introduced into the human HEK 293T cell line. We then delivered a second lentiviral vector encoding an Op*-specific gRNA (gRNA(Op*)) (or a non-specific gRNA (gRNA(NS)) as negative control) to these cells. Upon binding, gRNA(Op*) could mutate the critical dC residue in the WT Op repeat immediately upstream of its binding site, thus converting the Op repeat to a new Op* sequence that could serve as a new binding site for the same gRNA; this strategy enables sequential rounds of mutations (i.e., Op to Op* conversion) and gRNA binding events (Fig. 4A). We sequentially passaged cells harboring these circuits every three days for fifteen days (Fig. 4B) and observed GFP expression and the genotype of the cells by microscopy (Figs. 4C-D and S6A) and HTS (Figs. 4E-F), respectively. As shown in Fig. 4C, the frequency of GFP-positive cells in cultures harboring the 4xOp_1xOp*_GFP reporter and gRNA(Op*) increased over time, indicating the gradual activation of the reporter in the population. On the other hand, the frequency of GFP-positive cells did not change significantly in cultures that were transfected with gRNA(NS), or those that contained the 1xOp*_GFP reporter.

In addition to observing an increased frequency of GFP-positive cells, we observed that the intensity of the GFP signal in GFP-positive cells increased in cultures that harbored the 4xOp_1xOp*_GFP reporter and gRNA(Op*) over time (Fig. 4D). This data suggests that the number of bound transactivators, and thus, the number of activated (i.e., Op*) repeats that can serve as operator sites for the chimeric read-write-transactivator protein increased in these cells.

These results were further confirmed by analysis of the allele frequencies throughout the experiment by HTS. As shown in Fig. 4E, the frequency of the WT allele (state S0) in cells containing the repeat array and gRNA(Op*) decreased linearly with time over the course of the experiment. On the other hand, the frequency of intermediate states (S1 through S4) gradually increased and reached a plateau towards the end of the experiment, suggesting that these intermediate states reached steady state (Fig. 4F). The allele frequency of the final state (S5) gradually increased over the course of the experiment. No significant change in allele frequency was observed in cells that were transduced with a non-specific gRNA (Fig. S6B). Together with the microscopy data, these results show that the analog properties of a signal, such as the duration of exposure to gRNA(Op*), can be faithfully and permanently recorded within the distribution of memory states of the DNA recorder within the population. On the other hand, at the single cell level, each repeat forms a multi-bit digital recorder that associates longer or higher intensity of exposures to an incoming signal with transitions to higher memory states in the form of more accumulated mutations.

In samples harboring the gRNA(Op*) and either the 1xOp*_GFP or 4xOp_1xOP*_GFP reporters, we also observed dC to dG and dC to dA mutations, albeit with lower frequencies than for dC to dT mutations (Fig. S6C). This is consistent with previous results reported in mammalian cell lines (Komor et al., 2016; Nishida et al., 2016), and reflects the promiscuous outcome of repair of deaminated dC (dU) lesions in these cells. Notably, in samples containing the 1xOp*_GFP reporter, the frequency of the WT allele (state S0) decreased and the frequency of the mutant alleles increased linearly over time (Fig. S6C). Thus, even without having a repeat array, the accumulation of mutations in a specific target site can be used as an analog readout of an incoming signal.

In this experiment, we used VP64 as an activator domain. However, the activation level and dynamic range of the reporter output can be tuned by using stronger activator domains such as VPR (Chavez et al., 2015). Alternatively, other effector domains (such as repressors (Farzadfard et al., 2013), DNA methyl transferases (Liu et al., 2016), acetyl transferases (Hilton et al., 2015), or other types of histone modification domains) could be used to implement more sophisticated forms of gene regulation programs.

## Discussion

Our DOMINO platform addresses many limitations of current DNA writing platforms by using a DNA read-write head that converts the genomic DNA of living cells into a readable and writable medium that can be manipulated with single-nucleotide resolution. Orthogonal DOMINO operators can be built by simply changing the sequence of gRNAs, making the system highly scalable. Furthermore, due to the ability to manipulate DNA with single-nucleotide resolution within a defined window, compact multi-input operators can be readily created by targeting multiple gRNAs to nearby registers. By leveraging DNA as the computing and storage medium, we anticipate that this approach will be more stable than transcriptional memory strategies. Unlike other systems that require multiple recombinases to encode memory, DOMINO uses small gRNAs and only one protein moiety. The CRISPR-Cas9-based nature of this system and the absence of any requirement for double-strand DNA breaks or special repair mechanisms (such as Non-Homologous End Joining (NHEJ)) enables this system to be functional in both prokaryotic and eukaryotic cells. As a result, DOMINO offers a highly modular, robust and scalable strategy for dynamic programming of memory as well as order-independent, sequential and temporal logic operations in living cells. Furthermore, we show that DOMINO can be used to record both analog and digital signals, depending on the temporal nature of the circuits constructed. DOMINO circuits can be readily interfaced with other gene regulatory mechanisms to modulate gene expression and provide online readouts of cellular memory. Thus, we anticipate that DOMINO will allow for new strategies and unprecedented capacities to control cellular phenotypes and study biological phenomena in their native contexts.

In this paper, we focused on executing unidirectional DNA writing events by using a cytidine deaminase as the DNA writing module. Very recently, an adenosine deaminase DNA writing module that allows for dA to dG and dT to dC mutations was described (Gaudelli et al., 2017). Incorporating this new DNA writing module (or other orthogonal writer modules) into DOMINO should make reversible DNA writing possible, which has been challenging to achieve with previous DNA memory platforms. This will enable bidirectional cellular programs and thus pave the way for sophisticated biological state machines, cellular automata, and Turing machines that use the genomic DNA of living cells as a rewritable memory tape to perform advanced memory and computation operations.

In addition to digital computation, DOMINO operators can be used to perform analog memory and computation in living cells when propagation delays are taken into account. Furthermore, as shown in Figs. 3 and 4, analog properties (i.e. duration and magnitude) of an incoming signal can be recorded within the mutation states of the DOMINO operators. In these examples, recording capacity can be increased by extending the number of repeat elements or tuning the overlapping distance between the repeats. On the other hand, the input-output transfer function (i.e., the relationship between gRNA expression level and degree of mutation) can be tuned by adjusting the position of mutable residues within the gRNA WRITE window.

The self-reinforcing circuit presented in Fig. 4 can be used as the basis for building intelligent synthetic gene circuits with artificial learning capacities (Bray, 2003; Gandhi et al., 2007; Nesbeth et al., 2016; Tagkopoulos et al., 2008). Besides serving as a proof of concept for synthetic gene circuits with learning capacity, this circuit can be used as an online functional reporter for DNA memory states. Existing DNA-based molecular recording technologies rely on DNA sequencing as the readout. Thus, in these technologies, in order to retrieve the recorded information, the recording has to be stopped and cells need to be killed, which limit the applicability of these technologies to offline monitoring. On the other hand, the precise and sequential DNA writing achieved by DOMINO enables one to correlate the DNA memory state (i.e., the number of edited repeats) with the intensity of a fluorescence reporter signal that can be continuously monitored in living cells without disrupting the cells (Fig. 4A-D). This feature makes DOMINO recorders especially useful for studying biological events in an online fashion in their native context.

Deterministic DOMINO operators and cascades rely on precise base editing events for proper function. Our results show that using the CDA-nCas9-ugi head, the outcome of these operators in *E. coli* are almost exclusively in the form of dC to dT mutations. However, in human cells, other nucleotides (dG, and to a lesser extent, dA) are also generated, albeit with a lower rate than dT (Fig. S6C). In human cells, this issue could generate undesirable memory states that could reduce the performance of deterministic DOMINO operators. This can be addressed by implementing strategies that favor dC to dT mutations over the other possible outcomes to improve the efficiency of correct outcomes (Komor et al., 2017) or using alternative DNA writing modules that generate more pure editing products (Gaudelli et al., 2017).

Several CRISPR-Cas9 based strategies for recording information, such as signaling dynamics and cellular lineage histories, into DNA have been recently described (Frieda et al., 2017; Kalhor et al., 2016; McKenna et al., 2016). These approaches rely on stochastic DNA memory states (i.e., indel mutations) that are generated by Cas9-mediated double-strand DNA breaks and subsequent repair of these breaks by NHEJ. However, the recording capacity of these recorders are exhausted within a few generations or after recording a few molecular events due to loss of gRNA target sites and are therefore not ideal for long-term recording of signaling dynamics and event histories. Moreover, since indel mutations (memory states) are stochastically generated due to NHEJ, new mutations could destroy the previous mutations and thus overwrite the previous memory states, making tracing lineage histories challenging. In addition, none of these strategies can be used in organisms without an efficient NHEJ repair pathway, such as prokaryotes.

In contrast, mutational memory states generated by DOMINO are precise, unidirectional, position-specific, and minimally-disruptive. The features ensure that previous mutations are preserved after each editing step and can be accurately traced. The precise and predictable memory state transitions in DOMINO recorders enables one to couple memory states to functional biological outcomes, such as changes in gene expression (Fig. 4). Furthermore, DOMINO does not require double-strand DNA breaks or NHEJ, thus enabling it to function in both bacterial and mammalian cells in an autonomous and continuous fashion over many generations. We envision that the DNA record generated by the DOMINO recording system could be used to study signaling dynamics and event histories over many generations in their native contexts. The promiscuous repair of dC lesions in mammalian cells could actually be beneficial for lineage tracking applications, as it can increase the number of potential memory states. Moreover, signal-responsive lineage maps with tunable resolution can be generated because the activity of DOMINO recorder can be modulated by internal or external signals of interest. Combining these recorders with single-cell sequencing, advanced barcoding schemes, and self-targeting guide RNAs (Perli et al., 2016) should pave the way toward more advanced recorders for long-time monitoring of signaling dynamics and cellular lineages.

We envision that our long-term, compact, scalable, modular, and minimally disruptive DNA writers will enable an unprecedented set of applications for both building genetic programs and the recording of spatiotemporal molecular events in their native contexts. These applications could be highly impactful across many different fields, including development, cancer, stem cell differentiation, brain mapping, and many other areas. For example, DOMINO can be used to design and program the progression of developmental stages within living animals, or to perform long-term lineage tracking experiments in mammals, which has been impossible to date due to the lack of scalable and long-term methodologies. DOMINO recorders could be adapted to map neural activity by driving the activity of DNA writers with regulators that respond to neural activity. One could study the order and temporal nature of signaling events in their native contexts and robustly control cellular differentiation cascades *ex vivo* and *in vivo*. Our DNA writers could be programmed to investigate tumor development and unveil the cellular and environmental cues involved in tumor heterogeneity. Arbitrary information could be programmed into the DNA of living cells for DNA storage applications. Finally, living sensors could be designed to sense pathogens, toxins, or other signals within the body or in the environment and then later report on this information in detail. 18

## Acknowledgements

We thank Christina Harrison for helping with some of the early experiments in this project. This work was supported by the National Institutes of Health (P50 GM098792), the Office of Naval Research (N00014-13-1-0424), the National Science Foundation (MCB-1350625), the Defense Advanced Research Projects Agency, the MIT Center for Microbiome Informatics and Therapeutics, and NSF Expeditions in Computing Program Award 1522074.

## Supplementary Materials

Supplementary Figures 1-6

Supplementary Tables 1-3

Supplementary File 1

## Author contributions

F.F. conceived the study, designed and performed experiments and analyzed data. F.F. and N.G. designed the experiments, wrote the Sequalizer script, and analyzed next-generation sequencing data. F.F. and Y.H. performed the mammalian cell experiments and analyzed the results. G.J. and J.C. assisted with the bacterial experiments. T.K.L. supervised the research and provided scientific guidance and analysis. F.F., N.G., and T.K.L. wrote the manuscript with input from all authors.

## Supplementary Materials

### 1. Supplementary Text

#### Estimating Position-Specific Mutant Frequencies by Sequalizer

We developed a MATLAB program, dubbed Sequalizer (for <u>Seq</u>uence equ<u>alizer</u>), to calculate the frequency of base-pair substitutions in specific positions in a mixture of DNA species from Sanger sequencing chromatograms. Analyzing Sanger chromatograms by Sequalizer offers a low-cost strategy to HTS for assessing and quantifying frequency of precise mutations (i.e. nucleotide substitutions) that are generated by base-editing and other targeted genome engineering platforms.

Sequalizer uses a previously described algorithm (SeqDoC (Crowe, 2005)) to normalize and compute difference between Sanger chromatogram of a reference (unmodified) sequence and a test sample (which is expected to contain a mixture of DNA species containing mutations in specific positions). It then overlays the computed difference for all the four nucleotides (A, C, G, and T) on a single plot for the reference (top) and test sample (inverted, bottom) as a function of nucleotide position (x-axis) (Fig. S1A). A peak in this plot, indicates a difference in the normalized chromatogram signal between the reference and the test sample, and thus a mutation (i.e. base substitution) in that specific mutation. Sequlizer then estimates the frequency of mutants in each specific (targeted) position in the test sample using the difference between the heights of peaks corresponding to the reference and test samples in that position and reports that frequency as a number on top of the corresponding peaks. A test sample that has the same position-specific mutant frequency as the reference would result in no peaks in the Sequalizer plots (Fig. S1A, top panel). On the other hand, base-substitutions in the test sample compared to the reference sample can be detected as a peak in the Sequalizer plots (Fig. S1A, bottom panel). If a pure WT sample is used as the reference sample, the number printed on top of the peak estimates the frequency of molecules with mutation in that specific position in the test sample.

Since there is a high degree of variation between the height of peaks between different positions along a Sanger chromatogram, for each position Sequlizer normalizes the computed difference to the height of the peak for the reference chromatogram in that specific position. However, the height of the Sanger chromatogram containing 100% mutant alleles in a position could be different from the reference in that position, which could result in under- or overestimation of mutant frequencies by Sequalizer. Since the Sanger chromatogram, and thus the height of peaks for samples with the 100% mutant alleles are not always known, Sequlizer uses an experimentally determined parameter to account for the difference in height of peaks of Sanger chromatogram in each position. This parameter was calculated by mixing pure WT and pure mutant samples with different ratios, sequencing the mixtures, and using the Sequalizer output of the corresponding chromatograms to calculate a standard curve. As shown in Fig. S1B, the Sequalizer algorithm is able to compute frequencies of mutants at different positions solely based on Sanger chromatogram data, which correlates well with the mutant ratios in the mixtures.

We further verified Sequalizer by measuring position-specific mutant frequencies and comparing the output with the HTS for samples obtained from the order-independent AND gate circuit for the experiment described in Fig 1B. As shown in Fig. S1C, we observed high correlation (R^2^ values) between mutant frequencies measured by both methods in all the targeted positions, indicating that Sequalizer output can be used as a low-cost alternative to HTS. Deviation of the regression slope from unity (e.g., for C20 position) could be partially due to variations in the height of peaks of Sanger chromatograms between pure WT and pure mutant at different positions. As mentioned above, the Sequalizer algorithm tries to minimize the effect of such variations by normalizing the differences to the height of the WT peak in corresponding positions. However, since the heights of Sanger chromatograms for a pure mutant species also could affect the Sequalizer and this value is often unknown, it could cause the Sequalizer to underestimate or overestimate mutant frequencies compared to those measured by HTS. Nevertheless, the high correlation between Sequalizer outputs and HTS results indicate that changes in Sequalizer output can be used as a quantitative measure of changes in allele frequencies in a given position, even if they are not used for absolute measurements. The MATLAB script for Sequalizer is provided in Supplementary File 1. 32

### 2. Materials and Methods

#### 2.1 Strains and Plasmids

Standard molecular biology and cloning techniques, including ligation, Gibson assembly (Gibson, 2011) and Golden Gate assembly (Engler and Marillonnet, 2014) were used to construct the plasmids. Chemically competent *E. coli* DH5α F’ *lacI^q^* (NEB) and E. cloni 10G (Lucigen) were used for cloning. MG1655 PRO strain (MG1655 strain that harbors PRO cassette (pZS4Int-*lacI*/*tetR*, Expressys) and expresses *lacI* and *tetR* at high levels) (Lutz and Bujard, 1997) was used for all the bacterial experiments. HEK 293T cells (ATCC CRL-11268) were purchased from and authenticated by ATCC and were used for mammalian cell experiments. Lists of plasmids, synthetic parts and sequencing primers used in this study are provided in Tables S1, S2 and S3, respectively. Plasmids and their corresponding maps will be available on Addgene.

#### 2.2 Antibiotics and Inducers

For bacterial selection, antibiotics were used at the following concentrations: Carbenicillin (Carb, 50 μg/mL), and Chloramphenicol (Cam, 25-30 μg/mL).

For the experiments shown in Figs. 1E, 2D, 2E, S2C, and S4 different combinations of 200 ng/ml anhydrotetracycline (aTc), 0.1 mM Isopropyl β-D-1-thiogalactopyranoside (IPTG) and 0.2% Arabinose (Ara) were used to induce the corresponding circuits. For the experiments shown in Figs. S3 and S5, 250 ng/ml aTc and 0.005% Ara were used. For the experiment shown in Fig. 2A, 150 ng/ml aTc, 0.1 mM IPTG, and 0.2% Ara were used. For all the other experiments, unless otherwise noted, 250 ng/ml aTc, 1 mM IPTG and 0.2% Ara were used. All concentrations are final concentrations.

#### 2.3 Experimental procedure

##### 2.3.1 Bacterial Cell Experiments

Different plasmids expressing gRNAs and targets (listed in Table S1) were transformed into the reporter cells (MG1655 PRO) harboring aTc-inducible CDA-nCas9-ugi (for bacterial experiments, APOBEC1 CDA (Komor et al., 2016) was used as the writing module). Single transformant colonies were grown in LB + Carb + Cam for 6-8 hours to obtain seed cultures. Seed cultures were diluted (1:100) in fresh media containing different combinations of inducers and grown in 96-well plates for multiple days with serial dilution as indicated in induction patterns in corresponding figures. Samples for various analyses including HTS, Sequalizer, and flow cytometry were taken at indicated time points.

##### 2.3.2 Cell Cultures and Mammalian Cell Experiments

Cell culture and transfections were performed as described previously (3). HEK 293T cells were grown in DMEM supplemented with 10% fetal bovine serum (FBS) and 1% penicillin-streptomycin. Lentiviruses were packaged using the FUGW backbone (Addgene #25870) and psPAX2 and pVSV-G helper plasmids in HEK 293T cells. Filtered lentiviruses were used to infect respective cell lines in the presence of polybrene (8 μg/mL). Successful lentiviral integration was confirmed by using lentiviral plasmid constructs constitutively expressing fluorescent proteins or antibiotic resistance genes to serve as infection markers.

A lentiviral plasmid construct was made by placing the nCas9-CDA-ugi-VP64 fusion protein with nuclear localization signals linked to the Puromycin resistance gene with the P2A sequence under the control of constitutive CMV promoter (for mammalian experiments, PmCDA (Nishida et al., 2016) was used as the writing module). In addition, repeat arrays (4xOp_1xOp* or 1xOp*) were placed upstream of the minimal pMLV promoter driving EGFP and the resultant reporter constructs were cloned into the same lentiviral construct. The clonal cell lines harboring the two transcriptional units were constructed by infecting early passage HEK 293T cells with high titer lentiviral particles, selecting for pooled populations grown in the presence of Puromycin (7 µg/mL) and picking up clonal populations after seeding pooled population with the density of 0.5 cells per well in a 96-well plate.

On day 0, 440,000 clonal reporter cells were infected with high titer lentiviral particles encoding the sgRNAs driven by the U6 promoter in a 6-well plate with triplicates. Infection efficiency was more than 90% in every sample. The cells were harvested every 3 days until day 15 after the infection. Half of the harvested cells were seeded in a 6-well plate for further culture and a quarter of cells were collected for next-generation sequencing. Microscopic images were obtained just before the harvests.

##### 2.3.3 Microscopy Image Analysis

Fluorescence microscopy images of cells in tissue culture plates were obtained by using the ZEISS ZEN microscope software. For each sample, total number of EGFP-positive cells and signal intensities were measured from microscopic images of 5 random fields using CellProfiler image analysis software by using the ‘ColorToGray’, ‘IdentifyPrimaryObjects’, MeasureObjectIntensity’ and ‘ExportToSpreadsheet’ modules.

##### 2.3.4 Flow Cytometry

An LSR Fortessa II flow cytometer (Becton Dickinson, NJ) was used for all the experiments. GFP expression was measured using 488/FITC laser/filter set. All samples were uniformly gated and flow cytometry data were analyzed by FACSDiva and FlowJo (Becton Dickinson, NJ). For each gated sample, the mean fluorescence and percent of GFP-positive cells were calculated.

##### 2.3.5 High-throughput Sequencing

For each sample, 5 μl of culture was resuspended in 15 μl of QuickExtract DNA Extraction Solution (Epicentre, WI) and lysed by a two-step protocol (15 minutes incubation at 65 °C followed by 2 minutes incubation at 98 °C). Target sites were PCR amplified using 2 μl of lysed cultures as template and the appropriate primers listed in Table S3. The obtained amplicons were directly used as templates in a second round of PCR to add Illumina barcodes and adaptors. The amplicons were then multiplexed and analyzed by Illumina MiSeq. The obtained sequencing reads were demultiplexed and allele frequencies were calculated using a custom MATLAB script.

##### 2.3.6 Sanger Sequencing and Sequalizer Analysis

For each sample, target sites were PCR amplified by target-specific primers and Sanger sequenced by Quintara Biosciences. The obtained Sanger chromatograms were then analyzed by Sequalizer using seed cultures as reference as described above.

**Figure S1.**
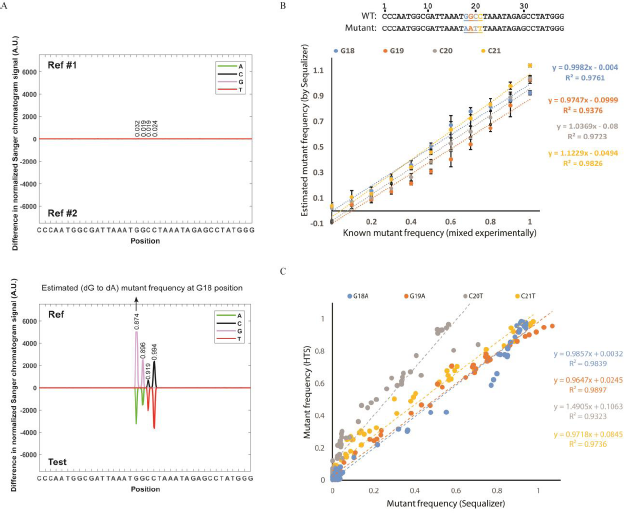
Using Sequalizer to estimate position-specific mutant frequencies from Sanger chromatograms. **A)** Sequalizer analysis comparing two instances of WT unmutated (i.e., Ref samples) sequences (top) and a WT unmutated (Ref) sequence vs. Test sample containing a mixture of mutated and unmutated sequences (bottom). The y-axis shows differences between normalized Sanger chromatograms for the samples being compared (Ref #1 vs. Ref #2 or Ref vs. Test). Peaks in these plots indicate differences in the normalized chromatograms and thus mutations in corresponding positions. For example, the peak marked by a black arrow in the bottom plot indicates mutations of dG at position 18 in the Ref to dA in the Test sample. The numbers above target positions (i.e., positions 18-21), show the estimated mutant frequency in that position based on the Sequalizer algorithm, which takes into account the height of Sanger chromatograms in a given position to normalize the calculated difference values. **B)** Standard curves obtained by analyzing samples containing known mutant ratios by Sequalizer. Two plasmids encoding the pure WT and mutant sequences (as indicated) were mixed at the following mutant:WT ratios: 0:10, 1:9, 2:8, 3:7, 4:6, 5:5, 6:4, 7:3, 8:2, 9:1, and 10:0. The mixtures were Sanger-sequenced and the obtained chromatograms were analyzed by Sequalizer. The estimated mutant frequencies at the four target positions were plotted against the known (i.e., experimentally mixed) mutant ratios. Error bars indicate standard deviation for six independent replicates. **C)** The position-specific mutant frequencies measured by Sequalizer vs. HTS at four target positions for samples from the experiment described in Fig. 1B.

**Figure S2.**
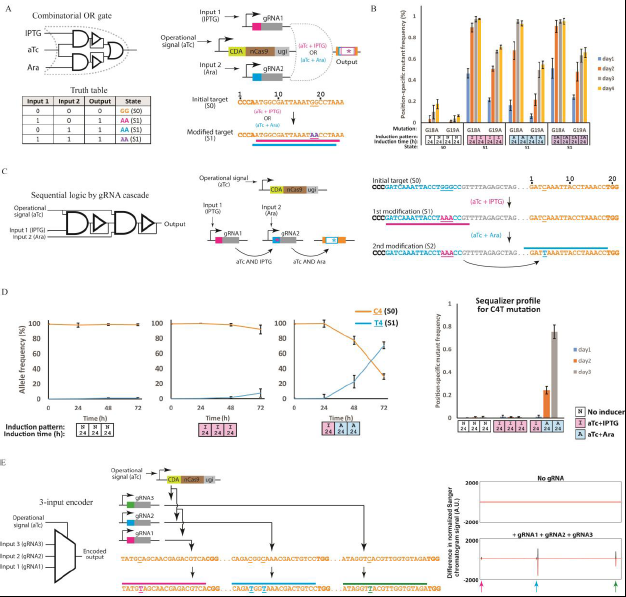
Examples of additional circuits built using DOMINO operators. **A)** Schematic representation and truth table for a DOMINO OR gate. **B)** Sequalizer results for the circuit shown in (A). *E. coli* cells were induced for four days using the indicated patterns and position-specific mutant frequencies were assessed by Sequalizer analysis of Sanger chromatograms. Error bars indicate standard deviation for three biological replicates. **C)** A sequential AND gate built by a cascade of gRNAs, where the first (IPTG-inducible) gRNA edits and activates a downstream gRNA, which can then edit a downstream target. As demonstrated in this example, gRNA outputs of a DOMINO cascade can be independently regulated by using inducible promoters, such as an Ara-inducible promoter. This offers greater flexibility compared to using mutations as DOMINO outputs (e.g., designs shown in Fig. 2 and 3). **D)** Dynamics of allele frequencies (i.e., memory states) for the circuit shown in (C) assessed by HTS (left panel) and population-averaged C4T mutation frequency assessed by Sequalizer (right panel). Error bars indicate standard deviation for three biological replicates. **E)** A multi-input encoder circuit, where the presence of three input gRNAs is converted into *cis*-encoded mutations in the same DNA target locus (*lacZ* ORF in *E. coli* in this case). The circuit can be used to encode multiple transcriptional signals from various loci across a genome into DNA memory within a confined region. The multiplexed and *cis*-encoded signals can then be read and decoded by HTS or Sanger sequencing to reveal information about the original signals. The plots on the right show the Sequalizer output for cells containing no gRNA (top) and those containing three constitutively expressed input gRNAs (bottom). Mutations in gRNA target sites are reflected as peaks in the bottom Sequalizer plot. This circuit is an example of a DOMINO circuit with more than two inputs, which we envision can be readily extended to additional inputs for *in vivo* memory applications and storing information (spatial, temporal, or artificial) across a genome.

**Figure S3.**
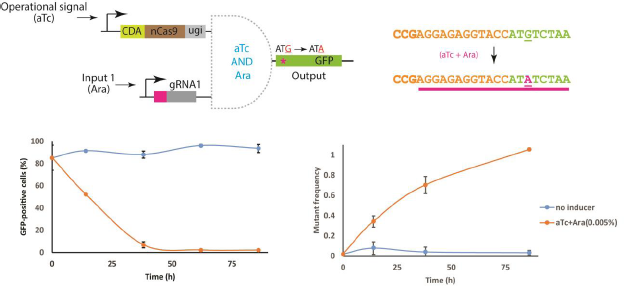
Regulation of gene expression by manipulating functional elements by DOMINO. Conditional conversion of a canonical, efficient initiation codon (ATG) to ATA (which is a non-efficient initiation codon) by an Ara-inducible DOMINO operator was used to down-regulate GFP expression in *E. coli*. Over time, the number of GFP-positive cells decreased and the frequency of mutants increased in induced samples while these quantities minimally changed in non-induced samples. For GFP measurements, samples were grown for six hours in LB with no inducers before flow cytometry to ensure removal of any repression (i.e., CRISPRi) effect enacted by bound CDA-nCas9-ugi. Error bars indicate standard deviation of three biological replicates.

**Figure S4.**
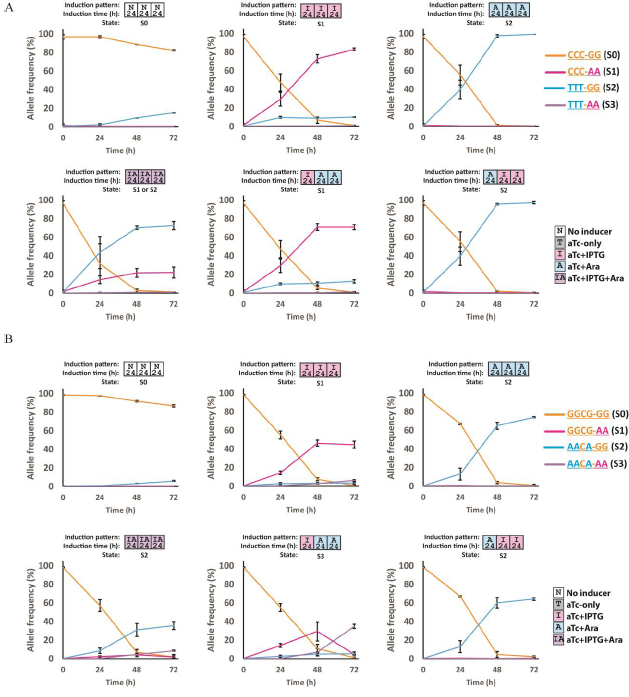
Dynamics of allele frequencies (memory states) for (A) the race-detecting circuit (Fig. 2D) and (B) the sequential logic circuit shown in Fig. 2E. In each subplot, the dominant allele in the last time point has been used to determine the memory state. Error bars indicate standard deviation for three biological replicates.

**Figure S5.**
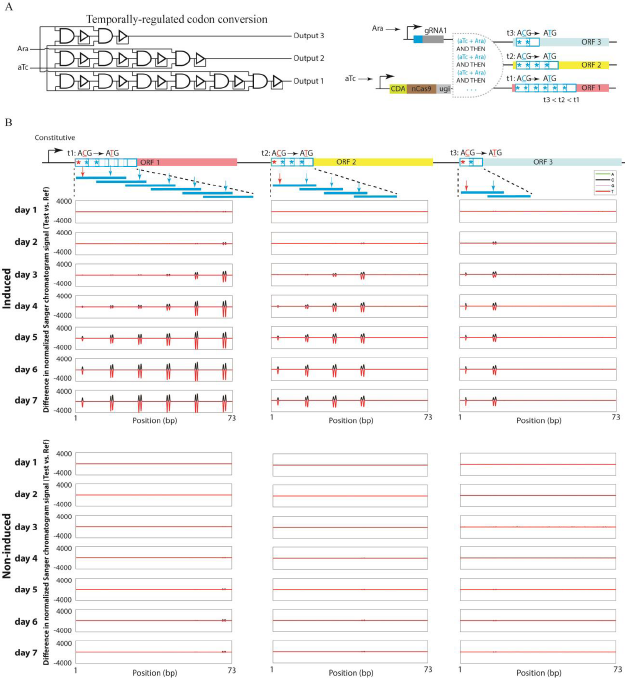
Using DOMINO delay elements to temporally control the conversion of cryptic start codons into canonical start codons in three ORFs. **A)** The schematic representation of the time-dependent codon conversion experiment. Three different ORFs with non-canonical (ACG) start codons and different number of delay elements (i.e., overlapping repeats) in their N-termini were placed in a synthetic operon. A gRNA was designed so that it could bind to the 3’-distal repeat element in each array. Sequential recruitment and editing of the repeat elements by this gRNA led to progressive mutation accumulation within the repeat elements toward the 5’-end and eventually editing of the upstream ACG codons to ATG. In this circuit, due to the presence of different number of delay elements in each array, different delay times and thus temporal regulation is achieved. The time required for start codon conversion for ORF 1 (t1) is expected to be longer than the time required for ORF 2 (t2) which itself is expected to be longer than the time required for the conversion in ORF 3 (t3). **B)** *E. coli* cells harboring the indicated circuit in (A) were induced and then mutation accumulation in the arrays was monitored by Sanger sequencing and Sequalizer over time. Upon induction of the circuit, time-dependent accumulation of mutations was observed in all the three repeat arrays. The position corresponding to the start codon (shown by red arrow) in the third ORF, which possessed only two repeats in its N-terminus array, was the first that accumulated significant levels of mutations. This was followed by the second ORF, which contained four delay elements and thus experienced a longer delay compared to ORF 3. The first ORF, which possessed six repeats and was thus subject to the longest delay, was the last ORF in which mutations in the position corresponding to the cryptic start codon were accumulated. On the other hand, in non-induced cells, only low levels of mutations accumulated in the downstream repeat of each array and only at the later time points of the experiment, likely due to the background activity of the promoters. Nevertheless, no mutations were detected in positions corresponding to cryptic start codons in non-induced cells.

**Figure S6.**
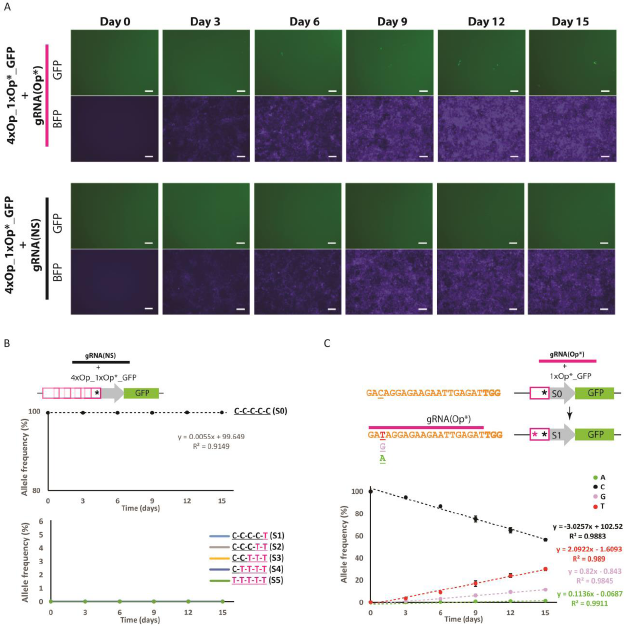
Representative microscopy images and additional data for the experiment shown in Fig. 4. **A)** Representative microscopy images for cells harboring the 4xOp_1xOp*_GFP reporter and the Op*-specific gRNA (gRNA(Op*)) or a non-specific gRNA (gRNA(NS)). Scale bars indicate 100 μm. **B)** Dynamics of allele frequencies (memory states) for cells harboring the 4xOp_1xOp*_GFP reporter and gRNA(NS) (negative control). **C)** Dynamics of allele frequencies (memory states) for cells harboring the 1xOp*_GFP reporter and gRNA(Op*). The mutable dC residue within the gRNA target site was mutated with a constant rate into dT and constant but lower rates into dG and dA, reflecting the promiscuous repair of deaminated cytidine lesions in mammalian cells. The linear decrease in dC allele frequency, as well as the linear increases in dT, dG, and dA allele frequencies, can be used as an analog readout of gRNA expression duration or intensity.

**Table S1.** **List of the plasmids used in this study**

| Name | Plasmid Code | Marker | Used in |
| --- | --- | --- | --- |
| P<sub>tetO</sub>_CDA-nCas9-ugi | pFF1454 | Cam | Figs. 1-3 & 5<br>Figs. S1-S5 |
| Comb_AND_gate | pFF1581 | Carb | Fig. 1B-D |
| Comb_AND_gate_gRNA_output | pFF1590 | Carb | Fig. 1E |
| Seq_AND_gate | pFF1610 | Carb | Fig. 2A-C |
| Race_detecting | pFF1684 | Cam | Fig. 2D<br>Fig. S4A |
| Mixed_seq_logic | pFF1685 | Carb | Fig. 2E<br>Fig. S4B |
| 3x_propagation_delay_seq_AND | pFF1588 | Carb | Fig. 3 |
| gRNA(Op*) | pYH383 | Carb<br>Hygro | Fig. 4<br>Fig. S6 |
| gRNA(NS) | pYH384 | Carb<br>Hygro | Fig. 4<br>Fig. S6 |
| 4xOp*_1xOp_GFP_pCMV_nCas9_CDA_ugi_VP64 | pYH396 | Carb<br>Puro | Fig. 4<br>Fig. S6 |
| 1xOp*_GFP_pCMV_nCas9_CDA_ugi_VP64 | pYH404 | Carb<br>Puro | Fig. 4<br>Fig. S6 |
| OR_gate | pFF1583 | Carb | Fig. S2A-B |
| gRNA_cascade | pFF1586 | Carb | Fig. S2C-D |
| Multiplexer | pFF1572 | Carb | Fig. S2E |
| Temporal_start_codon_conversion | pFF1573 | Carb | Fig. S5 |
| ATG_conversion | PFF1604 | Carb | Fig. S3 |

**Table S2.**
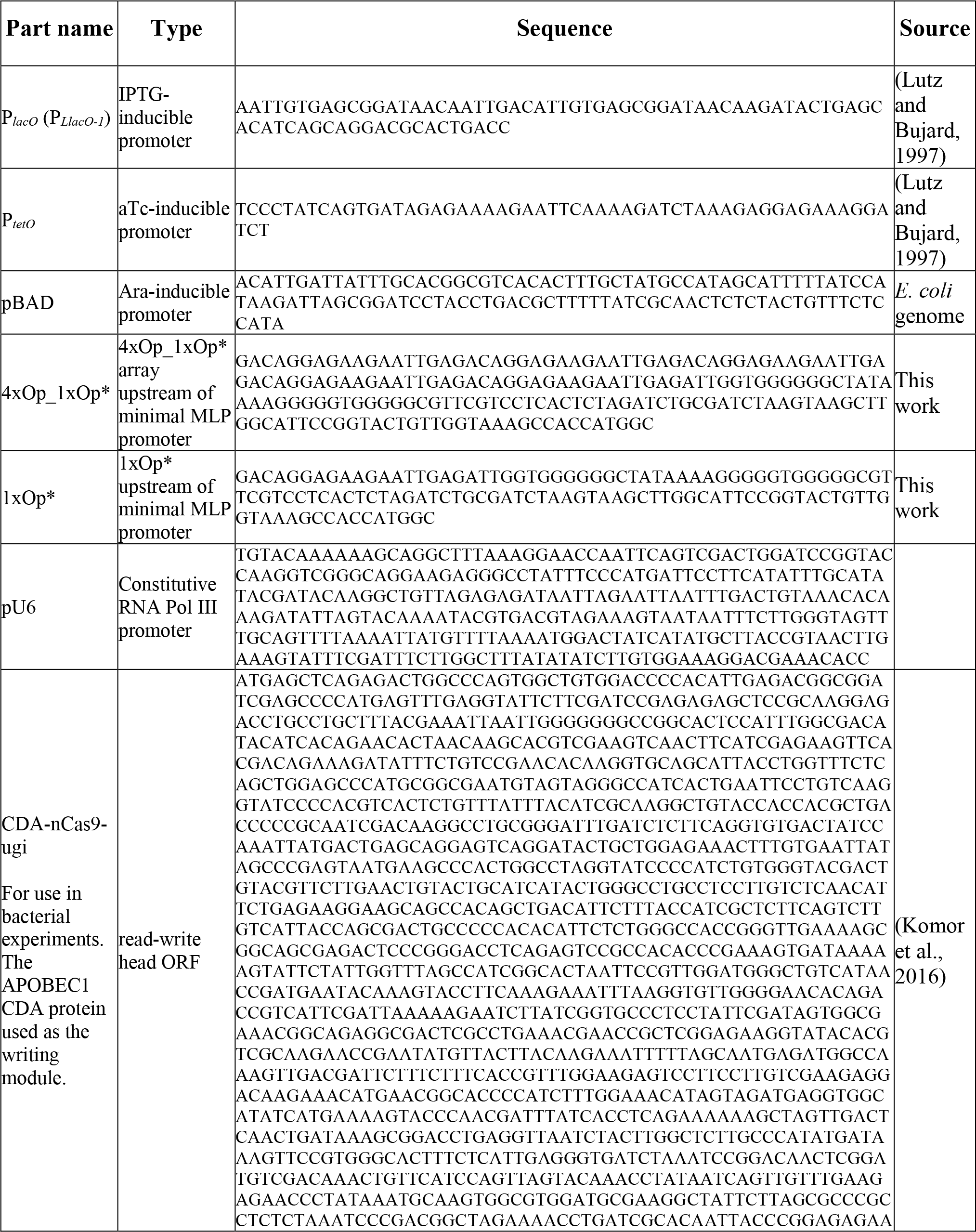
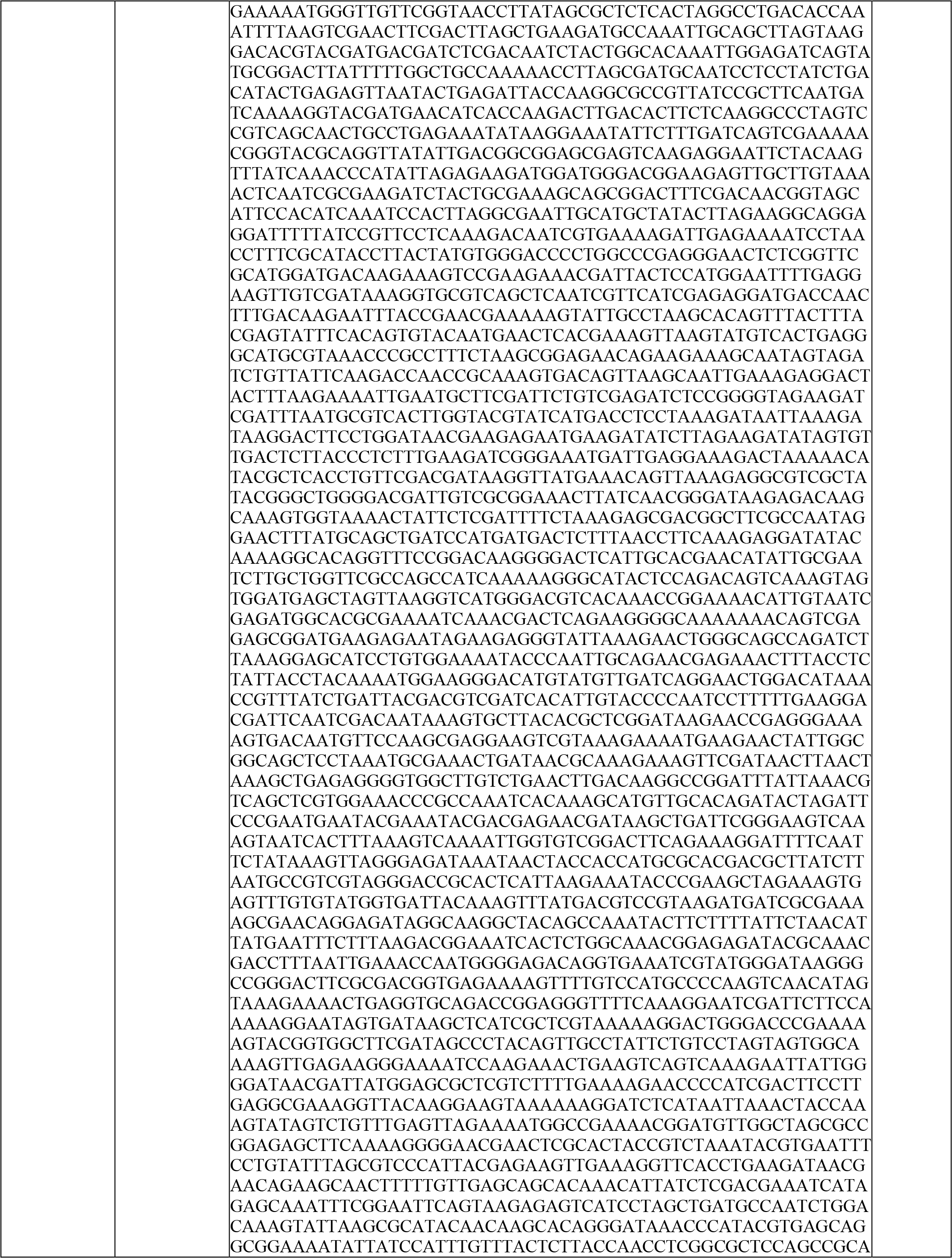
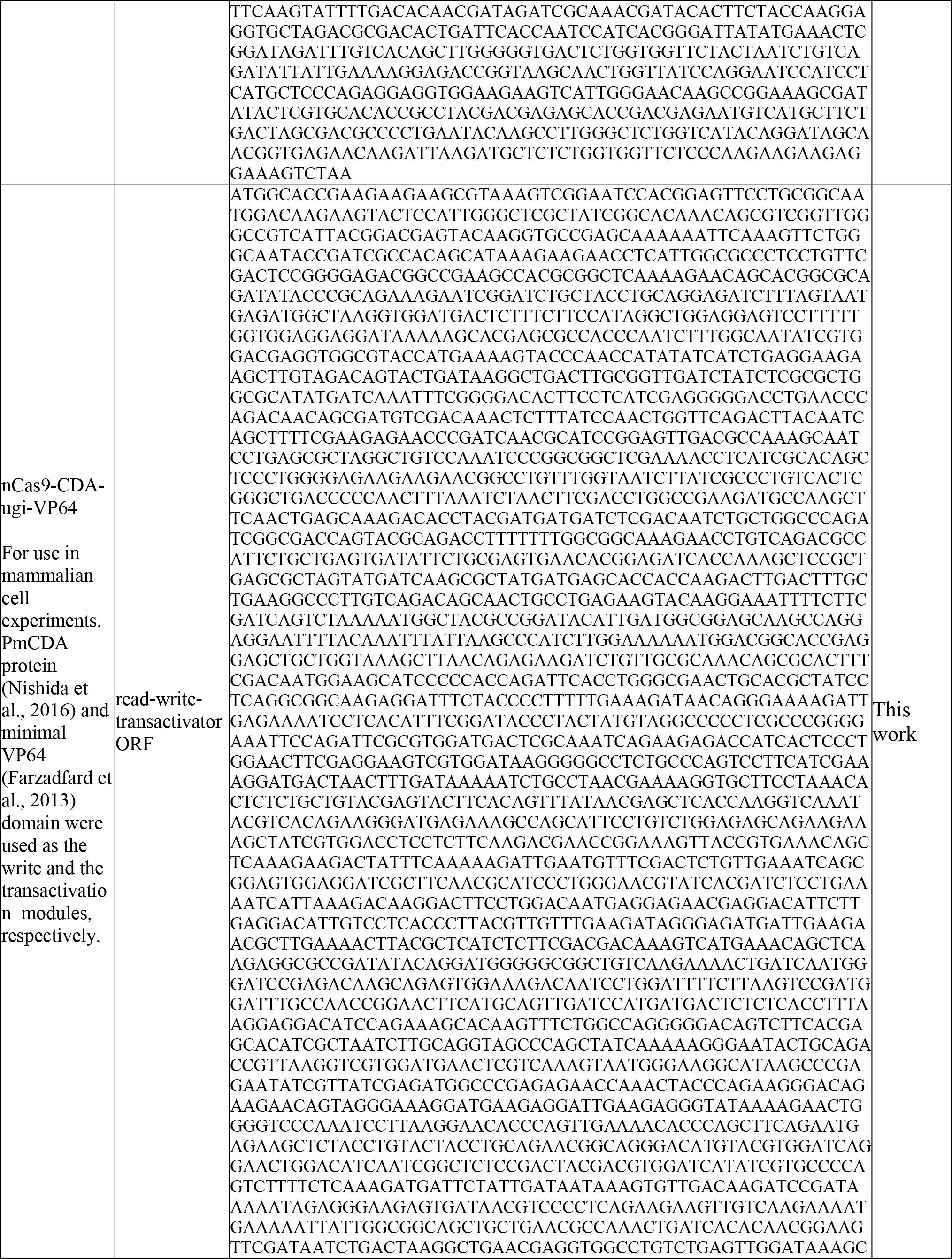
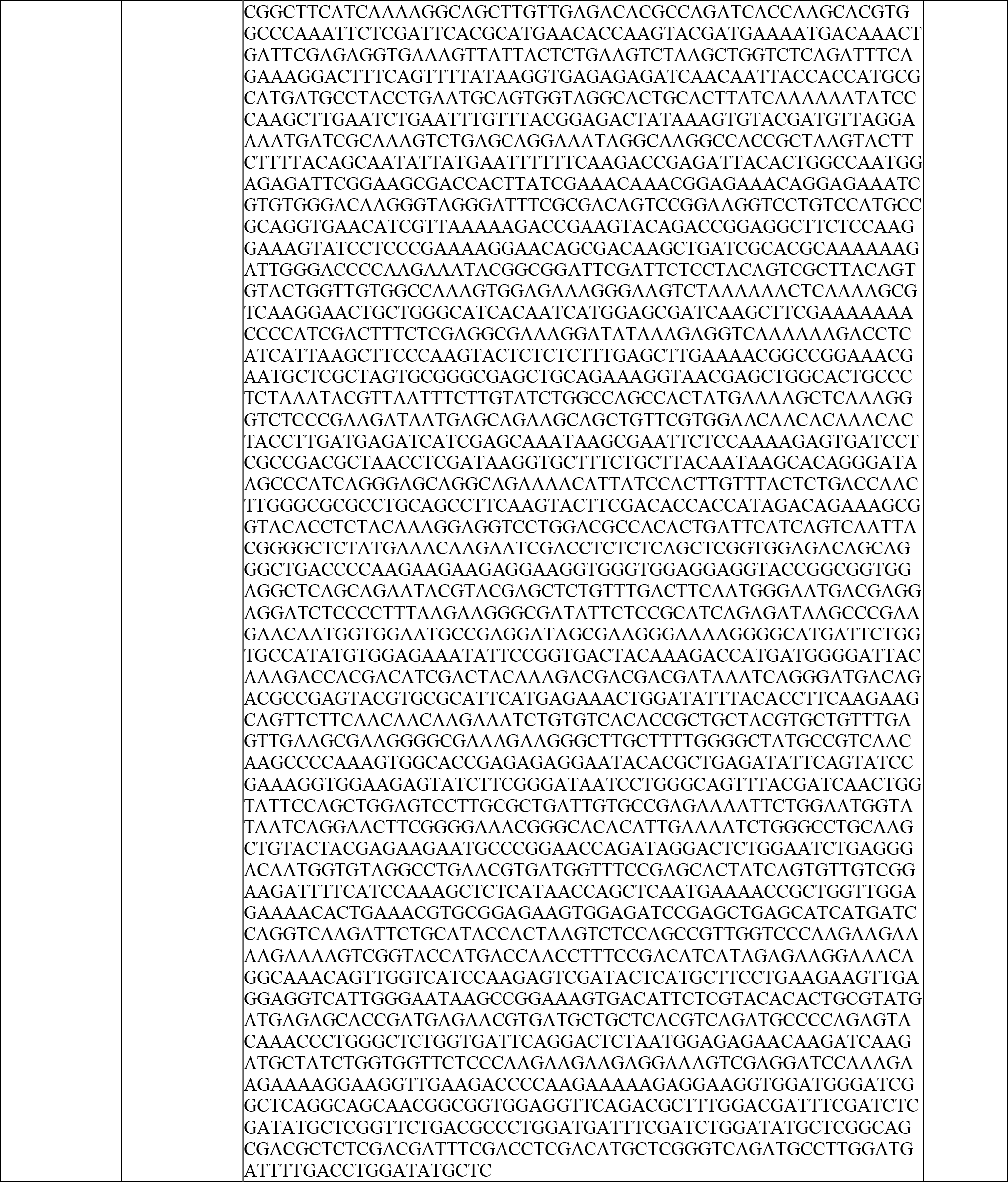
**List of the synthetic parts and their corresponding sequences used in this study**

**Table S3.**
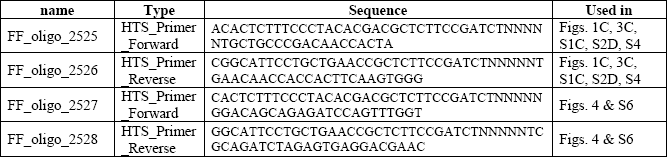
**List of HTS primers and their corresponding sequences used in this study**

## References and Notes

Arpino, J.A., Hancock, E.J., Anderson, J., Barahona, M., Stan, G.B., Papachristodoulou, A., and Polizzi, K. (2013). Tuning the dials of Synthetic Biology. Microbiology 159, 1236-1253.

Bray, D. (2003). Molecular networks: the top-down view. Science 301, 1864-1865.

Briner, A.E., Donohoue, P.D., Gomaa, A.A., Selle, K., Slorach, E.M., Nye, C.H., Haurwitz, R.E., Beisel, C.L., May, A.P., and Barrangou, R. (2014). Guide RNA functional modules direct Cas9 activity and orthogonality. Molecular cell 56, 333-339.

Chavez, A., Scheiman, J., Vora, S., Pruitt, B.W., Tuttle, M., E, P.R.I., Lin, S., Kiani, S., Guzman, C.D., Wiegand, D.J., et al. (2015). Highly efficient Cas9-mediated transcriptional programming. Nature methods 12, 326-328.

Coppoolse, E.R., de Vroomen, M.J., van Gennip, F., Hersmus, B.J., and van Haaren, M.J. (2005). Size does matter: cre-mediated somatic deletion efficiency depends on the distance between the target lox-sites. Plant molecular biology 58, 687-698.

Crowe, M.L. (2005). SeqDoC: rapid SNP and mutation detection by direct comparison of DNA sequence chromatograms. BMC bioinformatics 6, 133.

Engler, C., and Marillonnet, S. (2014). Golden Gate cloning. Methods in molecular biology 1116, 119-131.

Farzadfard, F., and Lu, T.K. (2014). Genomically encoded analog memory with precise in vivo DNA writing in living cell populations. Science 346, 1256272.

Farzadfard, F., Perli, S.D., and Lu, T.K. (2013). Tunable and multifunctional eukaryotic transcription factors based on CRISPR/Cas. ACS synthetic biology 2, 604-613.

Frieda, K.L., Linton, J.M., Hormoz, S., Choi, J., Chow, K.K., Singer, Z.S., Budde, M.W., Elowitz, M.B., and Cai, L. (2017). Synthetic recording and in situ readout of lineage information in single cells. Nature 541, 107-111.

Gandhi, N., Ashkenasy, G., and Tannenbaum, E. (2007). Associative learning in biochemical networks. Journal of theoretical biology 249, 58-66.

Gaudelli, N.M., Komor, A.C., Rees, H.A., Packer, M.S., Badran, A.H., Bryson, D.I., and Liu, D.R. (2017). Programmable base editing of A•T to G•C in genomic DNA without DNA cleavage. Nature advance online publication.

Gibson, D.G. (2011). Enzymatic assembly of overlapping DNA fragments. Methods in enzymology 498, 349-361.

Gilbert, L.A., Larson, M.H., Morsut, L., Liu, Z., Brar, G.A., Torres, S.E., Stern-Ginossar, N., Brandman, O., Whitehead, E.H., Doudna, J.A., et al. (2013). CRISPR-mediated modular RNA-guided regulation of transcription in eukaryotes. Cell 154, 442-451.

Hilton, I.B., D’Ippolito, A.M., Vockley, C.M., Thakore, P.I., Crawford, G.E., Reddy, T.E., and Gersbach, C.A. (2015). Epigenome editing by a CRISPR-Cas9-based acetyltransferase activates genes from promoters and enhancers. Nature biotechnology 33, 510-517.

Kalhor, R., Mali, P., and Church, G.M. (2016). Rapidly evolving homing CRISPR barcodes. Nature methods.

Kalhor, R., Mali, P., and Church, G.M. (2017). Rapidly evolving homing CRISPR barcodes. Nature methods 14, 195-200.

Komor, A.C., Kim, Y.B., Packer, M.S., Zuris, J.A., and Liu, D.R. (2016). Programmable editing of a target base in genomic DNA without double-stranded DNA cleavage. Nature 533, 420-424.

Komor, A.C., Zhao, K.T., Packer, M.S., Gaudelli, N.M., Waterbury, A.L., Koblan, L.W., Kim, Y.B., Badran, A.H., and Liu, D.R. (2017). Improved base excision repair inhibition and bacteriophage Mu Gam protein yields C:G-to-T:A base editors with higher efficiency and product purity. Science advances 3, eaao4774.

Lee, J.W., Gyorgy, A., Cameron, D.E., Pyenson, N., Choi, K.R., Way, J.C., Silver, P.A., Del Vecchio, D., and Collins, J.J. (2016). Creating Single-Copy Genetic Circuits. Molecular cell 63, 329-336.

Liu, X.S., Wu, H., Ji, X., Stelzer, Y., Wu, X., Czauderna, S., Shu, J., Dadon, D., Young, R.A., and Jaenisch, R. (2016). Editing DNA Methylation in the Mammalian Genome. Cell 167, 233-247 e217.

Lutz, R., and Bujard, H. (1997). Independent and tight regulation of transcriptional units in Escherichia coli via the LacR/O, the TetR/O and AraC/I1-I2 regulatory elements. Nucleic acids research 25, 1203-1210.

McKenna, A., Findlay, G.M., Gagnon, J.A., Horwitz, M.S., Schier, A.F., and Shendure, J. (2016). Whole-organism lineage tracing by combinatorial and cumulative genome editing. Science 353, aaf7907.

Nesbeth, D.N., Zaikin, A., Saka, Y., Romano, M.C., Giuraniuc, C.V., Kanakov, O., and Laptyeva, T. (2016). Synthetic biology routes to bio-artificial intelligence. Essays in biochemistry 60, 381-391.

Nishida, K., Arazoe, T., Yachie, N., Banno, S., Kakimoto, M., Tabata, M., Mochizuki, M., Miyabe, A., Araki, M., Hara, K.Y., et al. (2016). Targeted nucleotide editing using hybrid prokaryotic and vertebrate adaptive immune systems. Science 353.

Perli, S.D., Cui, C.H., and Lu, T.K. (2016). Continuous genetic recording with self-targeting CRISPR-Cas in human cells. Science 353.

Qi, L.S., Larson, M.H., Gilbert, L.A., Doudna, J.A., Weissman, J.S., Arkin, A.P., and Lim, W.A. (2013). Repurposing CRISPR as an RNA-guided platform for sequence-specific control of gene expression. Cell 152, 1173-1183.

Roquet, N., Soleimany, A.P., Ferris, A.C., Aaronson, S., and Lu, T.K. (2016). Synthetic recombinase-based state machines in living cells. Science 353, aad8559.

Siuti, P., Yazbek, J., and Lu, T.K. (2013). Synthetic circuits integrating logic and memory in living cells. Nature Biotechnology 31, 448-452.

Stark, W.M. (2017). Making serine integrases work for us. Current opinion in microbiology 38, 130-136.

Tagkopoulos, I., Liu, Y.C., and Tavazoie, S. (2008). Predictive behavior within microbial genetic networks. Science 320, 1313-1317.

